# Structural Insights into the Iron Nitrogenase Complex

**DOI:** 10.1101/2023.05.02.539077

**Authors:** Frederik V. Schmidt, Luca Schulz, Jan Zarzycki, Niels N. Oehlmann, Simone Prinz, Tobias J. Erb, Johannes G. Rebelein

## Abstract

Nitrogenases are best known for catalysing the reduction of dinitrogen to ammonia at a complex metallic cofactor. Recently, nitrogenases were shown to reduce carbon dioxide (CO_2_) and carbon monoxide to hydrocarbons, offering a pathway to recycle carbon waste into hydrocarbon products. Among the nitrogenase family the iron nitrogenase is the isozyme with the highest wildtype activity for the reduction of CO_2_, but the molecular architecture facilitating these activities remained unknown. Here, we report a 2.35-Å cryogenic electron microscopy structure of the Fe nitrogenase complex from *Rhodobacter capsulatus,* revealing an [Fe_8_S_9_C*-(R*)-homocitrate]-cluster in the active site. The enzyme complex suggests that the AnfG-subunit is involved in cluster stabilisation, substrate channelling and confers specificity between nitrogenase reductase and catalytic components. Moreover, the structure highlights a different interface between the two catalytic halves of the iron and the molybdenum nitrogenase, potentially influencing the intra-subunit ‘communication’ and thus the nitrogenase mechanism.

## Introduction

Nitrogenases catalyse a key step in the global nitrogen cycle by reducing molecular nitrogen (N_2_) to ammonia (NH_3_). Together with the energy-intensive industrial Haber-Bosch process, nitrogenases provide the vast majority of bioavailable nitrogen, which is essential for all life on Earth to build central metabolites such as nucleotides and amino acids [1-3]. Due to the extraordinary ability of nitrogenases to break the stable N≡N triple bond under ambient conditions, the mechanism of nitrogenases has been under scrutiny for decades [4-8].

Three homologous nitrogenase isoforms are known to date [9, 10]. The most prevalent and best studied nitrogenase is the molybdenum (Mo) nitrogenase (encoded by *nifHDK*), which is present in all known diazotrophs. Some diazotrophs encode ‘back-up’ or alternative nitrogenase genes: *vnfHDGK* for the vanadium (V) or *anfHDGK* for the iron (Fe) nitrogenase, expressed upon the depletion of Mo. All three nitrogenases consist of two components, the reductase component (NifH_2_, VnfH_2_, AnfH_2_) and the catalytic component (Nif(DK)_2_, Vnf(DGK)_2_, Anf(DGK)_2_). Importantly, the catalytic component of both alternative nitrogenases contains an additional subunit (VnfG or AnfG), whose role and function remains elusive.

The homodimeric reductase component contains a [Fe_4_S_4_]-cluster and two adenosine triphosphate (ATP) binding sites. In the ATP-bound state the reductase component transiently associates with its catalytic component. Upon complex formation, low-potential electrons are transferred from the [Fe_4_S_4_]-cluster of the reductase component via an [Fe_8_S_7_]-relay (P-cluster) to the active site cofactor of the catalytic component. The active site cofactor follows the general composition of [MFe_7_S_9_C-(*R*)-homocitrate], where M is either Mo, V, or Fe, depending on the nitrogenase isoform. Based on the containing heterometal, the clusters are termed FeMoco, FeVco or FeFeco. The structures of the FeMoco, the FeVco and just recently the FeFeco have been structurally confirmed by X-Ray crystallography [11, 12]. These studies revealed that in the FeVco one of the belt-sulphur (S) atoms is replaced by a carbonate, resulting in a [VFe_7_S_8_C(CO_3_)^2-^)(*R*)-homocitrate]-cluster. Based on the similar architecture of nitrogenases, one might expect them to follow one general catalytic mechanism. However, under 1 atm N_2_ and high electron flux conditions (ratio of reductase to catalytic component ≥ 20) different amounts of H_2_ are produced per mole N_2_ reduced (equations 1-3) [13].

Mo nitrogenase:

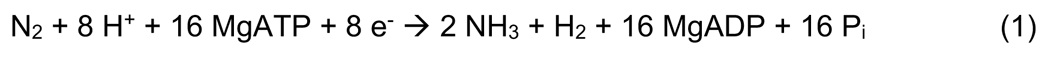

V nitrogenase:

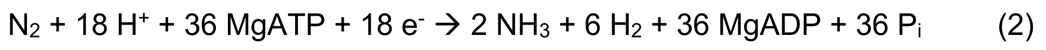

Fe nitrogenase:

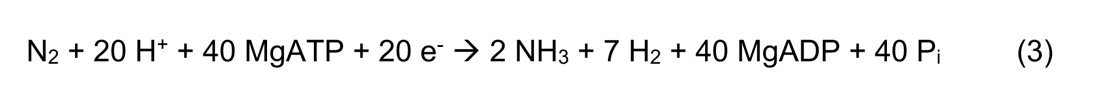

Lately, it was discovered that besides N_2_ all three nitrogenases can also reduce carbon monoxide (CO) to hydrocarbons. For CO reduction, the V nitrogenase is the most active isoform, mainly forming C-C bonds and releasing C_1_ to C_4_ hydrocarbons, mainly ethylene [14, 15]. The Fe nitrogenase shows around one third of the CO-activity compared to the V nitrogenase but only releases methane [16]. The Mo nitrogenase converts CO exclusively into C_2_ to C_4_ hydrocarbon chains but is ∼800-fold less active than the V nitrogenase [17]. This CO-processing activity can also be exploited for the *in vivo* conversion of the industrial exhaust CO to hydrocarbons as demonstrated for *Azotobacter vinelandii* expressing the V nitrogenase [18].

Beyond CO, it was recently shown that the wildtype V nitrogenase also reduces carbon dioxide (CO_2_) to CO, ethene and ethane [19]. In contrast, the wildtype Mo nitrogenase reduces CO_2_ only to CO [20] and formate [21]. Surprisingly, the Fe nitrogenase shows the highest CO_2_ reduction activity among the wildtype nitrogenases, converting CO_2_ to methane and formate [22]. The tremendous activity differences and varying product spectra, particularly for the reduction of CO and CO_2_ (further reviewed here: [23, 24]), suggest distinct differences among the three nitrogenase isoenzymes, which are not yet fully understood.

To gain molecular insights into the differences among the three nitrogenase isoenzymes, we set out to solve the structure of the Fe nitrogenase complex. For this, we expressed, purified and characterised the Fe nitrogenase of the phototroph *Rhodobacter capsulatus* in its native host. Using anaerobic single-particle cryogenic electron microscopy (cryoEM), we solved the structure of the adenosine diphosphate-aluminium fluoride (ADP-AlF_3_)-stabilized Fe nitrogenase complex consisting of two reductase and one catalytic component at a resolution of 2.35 Å. The structure of the Fe nitrogenase reveals the molecular architecture of the FeFeco and suggests three potential roles of the G-subunit: *i*) binding of the reductase component, *ii*) substrate channelling and *iii*) FeFeco positioning and stabilization. Furthermore, the structure allows us to compare the entire Fe nitrogenase complex with previously published Mo nitrogenase complexes [25-27] and the catalytic component of the V nitrogenase [12]. The comparison reveals distinct features of the Fe nitrogenase architecture, which distinguishes it from the Mo nitrogenase and might influence the catalytic mechanism of the alternative nitrogenases.

## Results

### Engineering *Rhodobacter capsulatus* for nitrogenase expression

We engineered *R. capsulatus* for studying the Fe nitrogenase. The purple non-sulphur bacterium *R. capsulatus* naturally harbours the Mo and Fe nitrogenase, its genome has been fully sequenced [28] and basic molecular biology methods have been established [29]. Using the *sacB* scar-less deletion system (see materials & methods), we engineered *R. capsulatus* B10S [30] to enable high-yield recombinant production and purification of the Fe nitrogenase. (*i)* We deleted the Mo nitrogenase-encoding gene cluster (*nifHDK*) to ensure that only the alternative nitrogenase is expressed. (*ii)* We knocked out the high-affinity molybdenum transporter genes *modABC* [31]. This modification is essential for high expression levels of the Fe nitrogenase, since trace amounts of molybdenum inside the cell repress the transcription of the Fe nitrogenase genes. (*iii)* We deleted a post-translational modification mechanism that inactivates the nitrogenase reductase component through ADP-ribosylation, encoded by *draT* and *draG* [32]. (*iv)* We removed the bacterial capsule by deleting *gtaI*. Previously, this knockout was found to improve the cell pellet quality post centrifugation [33], thus rendering this modification particularly useful for large-scale protein purification with *R. capsulatus*. (*v)* We interrupted the Fe nitrogenase locus *anfHDGK* by introducing a gentamycin resistance cassette, which allows the recombinant production of the affinity-tagged Fe nitrogenase from expression plasmids. For this purpose, we cloned the *anfHDGK* operon from the bacterial genome into a pOGG024-*kanR* vector and fused a His_6_-tag to the AnfH N-terminus and a Strep-tag II to the C-terminus of AnfD. For nitrogenase expression, we used conjugation to transfer the plasmid into the modified *R. capsulatus* strain. All genetic modifications of the final strain were confirmed by next generation sequencing (Table 1, Extended Data Fig. 1). In summary, we introduced five genetic modifications into *R. capsulatus* that render the purple non-sulphur bacterium an ideal platform for the plasmid-based production and characterisation of the Fe nitrogenase. The plasmid-based expression of nitrogenases in *R. capsulatus* complements the chromosomal nitrogenase expression of *A. vinelandii*, which has been the standard in the field so far.

**Fig. 1:**
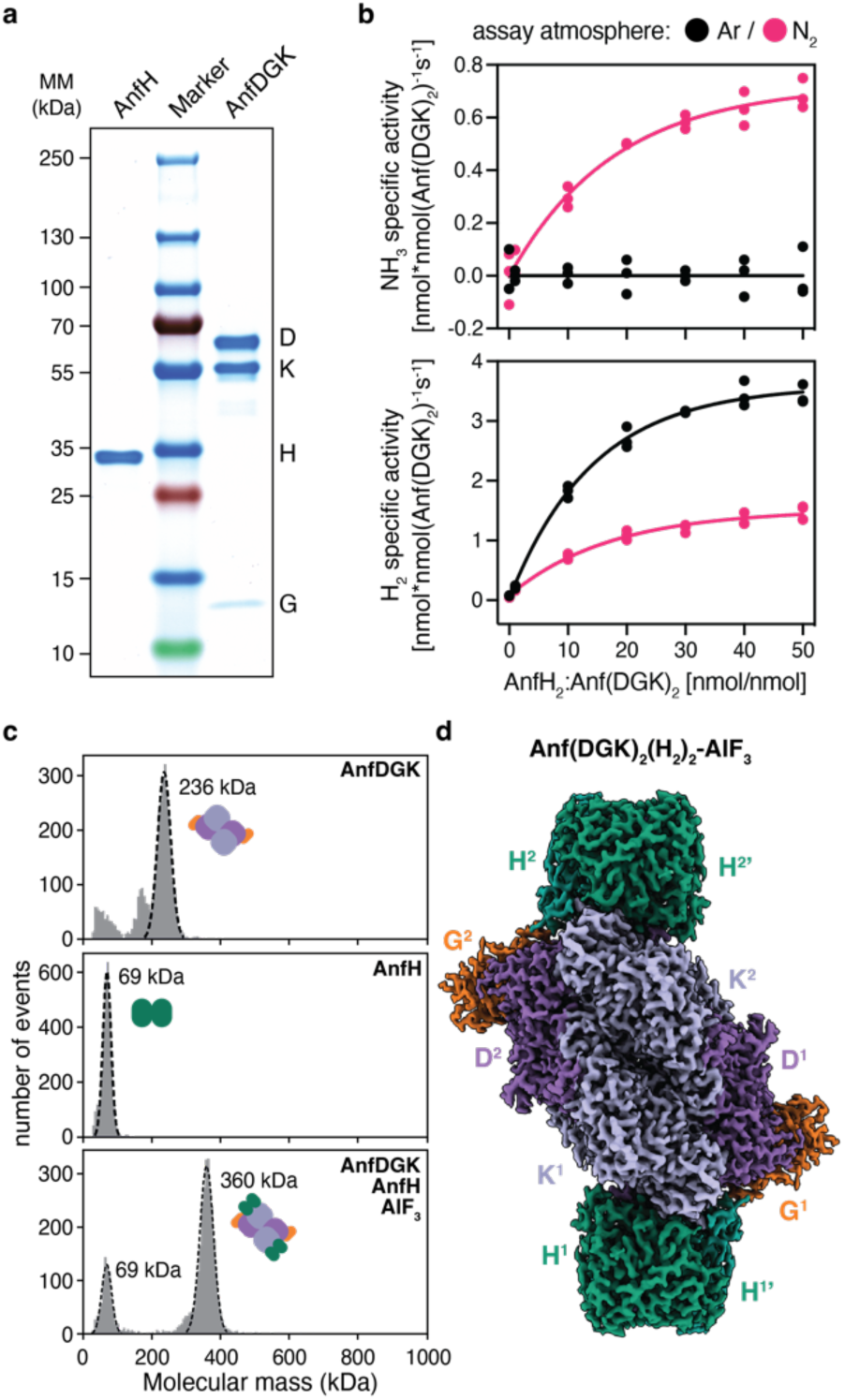
Purification and biochemical characterisation of the Fe nitrogenase. (**a**) SDS-PAGE analysis of the purified Fe nitrogenase reductase component (AnfH) and catalytic component (AnfDGK). (**b**) *In vitro* activity assays of the purified Fe nitrogenase under Ar or N2 atmosphere. Plotted are the specific activities for NH3 (top) or H2 (bottom) formation under varying molar ratios of AnfH2 to Anf(DGK)2. Individual measurements (n =3) are shown and the solid line represents the nonlinear fit of the data. (**c**) Mass photometry analysis of the individual nitrogenase components (top and middle) and the AlF3-trapped Anf(DGK)2(H2)2 complex (bottom). Plotted are the number of events versus the molecular mass of the individual events (in kDa). (**d**) Electron density map of the AlF3-trapped Fe nitrogenase complex at a global resolution of 2.35 Å.

### Purification and *in vitro* characterisation of the Fe nitrogenase

Using our *R. capsulatus* expression strain, we purified and biochemically characterised the Fe nitrogenase. As described in the materials and methods section, we established an anaerobic workflow for the separate purification of the reductase and catalytic components (Fig. 1a). *In vitro*, the Fe nitrogenase converted dinitrogen (N_2_) to ammonia (NH_3_) at a maximal rate of 0.69 nmol × nmol (Anf(DGK)2)^-1^ × s^-1^, closely matching the previously published value of 0.72 nmol × nmol (Anf(DGK)2)^-1^ × s^-1^ [34]. Notably, the rate of dihydrogen (H_2_) formed under N_2_ is twice as high as the rate of NH_3_ formation. As expected, these rates were found to follow a hyperbolic trend with increasing product formation proportional to the ratio of reductase to catalytic component (Fig. 1b). In a pure argon (Ar) atmosphere all electrons are directed towards H_2_ formation and a maximal rate of 3.44 nmol × nmol (Anf(DGK)2) ^-1^ × s^-1^ was measured, approximately double the H_2_ formation rate of the N_2_ atmosphere (Fig. 1b). Metal quantification via inductively coupled plasma optical emission spectroscopy (ICP-OES) suggested full Fe occupancy for the reductase component and ∼80% occupancy for the catalytic component (Extended Data Fig. 2). This result might be caused by a partial cluster occupancy of the catalytic component or could be the result of slight impurities in the Anf(DGK)_2_ samples (Fig. 1a and 1c). However, no transition metal other than iron was detected in our protein samples, confirming a pure Fe nitrogenase. Next, we analysed the complex formation of the Fe nitrogenase *in vitro* by trapping the ADP-bound reductase component on the Anf(DGK)_2_ core with AlF_3_. Following size exclusion chromatography (SEC), we could detect a protein complex of ∼360 kDa in size (Fig. 1c, bottom). The measured masses of the individual nitrogenase components were 236 kDa for Anf(DGK)_2_ and 69 kDa for AnfH_2_ (Fig. 1c, top and middle), thus indicating an Anf(DGK)_2_(H_2_)_2_ stoichiometry of the complex. These results agree with analytical SEC (Fig. 2c and Extended Data Fig. 2). Next, we analysed the high molecular weight complex by cryoEM. Following anaerobic sample preparation including plunge freezing inside an anaerobic tent, we obtained a 2.35 Å map visualising the expected heterodecameric complex of two AnfH_2_ dimers bound to Anf(DGK)_2_ (Fig. 1d). Taken together, we purified a fully active Fe nitrogenase from *R. capsulatus*, analysed its activity and complex formation *in vitro* and solved the structure of the AnfH_2_-bound complex.

**Fig. 2:**
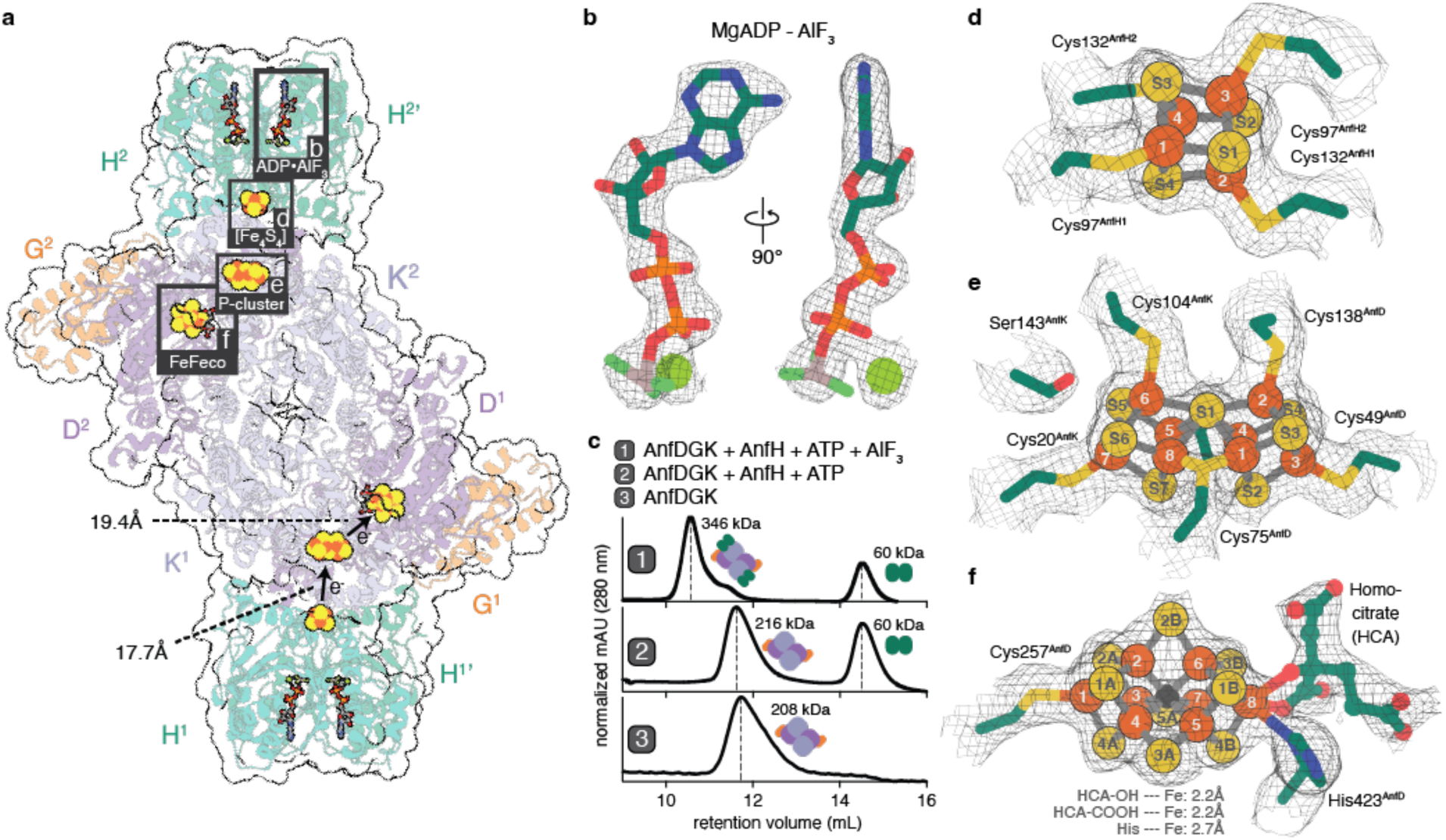
Structure of the Fe nitrogenase and its cofactors. (**a**) Model of the Fe nitrogenase fitted into the cryoEM electron density map. Reductase components (H^1^ + H^1^’ and H^2^ + H^2’^) are shown in green and the catalytic component in purple (D^1^ and D^2^), light purple (K^1^ and K^2^) and orange (G^1^ and G^2^). Cofactors are highlighted by grey boxes numbered b, d – e. Distances from the [Fe4S4]-to the P-cluster and from the P-cluster to FeFeco are indicated. (**b**) MgADP-AlF3 cofactor bound to AnfH. Carbon is coloured in green, nitrogen in blue, oxygen in red, phosphor in orange, aluminium in grey and fluoride in light green. The magnesium ion is depicted as a light green sphere. (**c**) Comparison between the elution profiles of a SEC performed with (1) AnfDGK and AnfH in the presence of ATP and AlF3, (2) AnfDGK and AnfH in the presence of ATP and (3) AnfDGK alone. The reductase component bound Fe nitrogenase complex only elutes in the presence of ATP and AlF3, reasoning our decision to model AlF3 instead of a γ-phosphate as shown in (b). (**d**) – (**f**) Close-up views on (d) the [Fe4S4]-cluster, (e) the P-cluster and the (f) FeFeco. Sulphur atoms of the clusters are represented as yellow spheres, the according iron atoms as orange spheres and the central carbide ion of the FeFeco as a black sphere. Amino acid residues in the direct cofactor environment are depicted as sticks, with carbon coloured in green, sulphur in yellow, nitrogen in blue and oxygen in red. The homocitrate coordinated to the FeFeco is depicted as a ball-stick model with the same colour coding as for the amino acid residues.

### Structure of the Fe nitrogenase

Using the cryoEM map, we created a model of the Fe nitrogenase complex and analysed its molecular features. We used AlphaFold [35] models for the catalytic component (Anf(DGK)_2_) and the previously published crystal structure of the reductase component (AnfH_2_) from *A. vindelandii* (PDB: 7QQA, [36]) to build a detailed model of the Fe nitrogenase into the electron density map (Fig. 2a). All nitrogenase cofactors were well resolved with local resolutions of up to 1.83 Å (Fig. 2b, d-f, Extended Data Fig. 3). To facilitate the electron transport between the [Fe_4_S_4_]-cluster of the reductase component to the P-cluster of the catalytic component, two molecules of MgATP must bind to AnfH_2_ forming a nucleotide-dependent Fe nitrogenase complex. Indeed, our structure contains one ATP mimic MgADP-AlF_3,_ per AnfH subunit, locking the Fe nitrogenase complex in the transition state (Fig. 2a and b) [37-39]. Although ATP also fits into the electron density of the cofactor, we decided to model AlF_3_ at the terminal end of the phosphate esters based on our observation that the AnfH_2_-bound complex only eluted in the presence of ATP and AlF_3_ during SEC (Fig. 2c). In the dimeric interface of the reductase component, we observed a [Fe_4_S_4_]-cluster coordinated by Cys97 and Cys132 of the two interacting AnfH subunits (Fig. 2a and d).

**Fig. 3:**
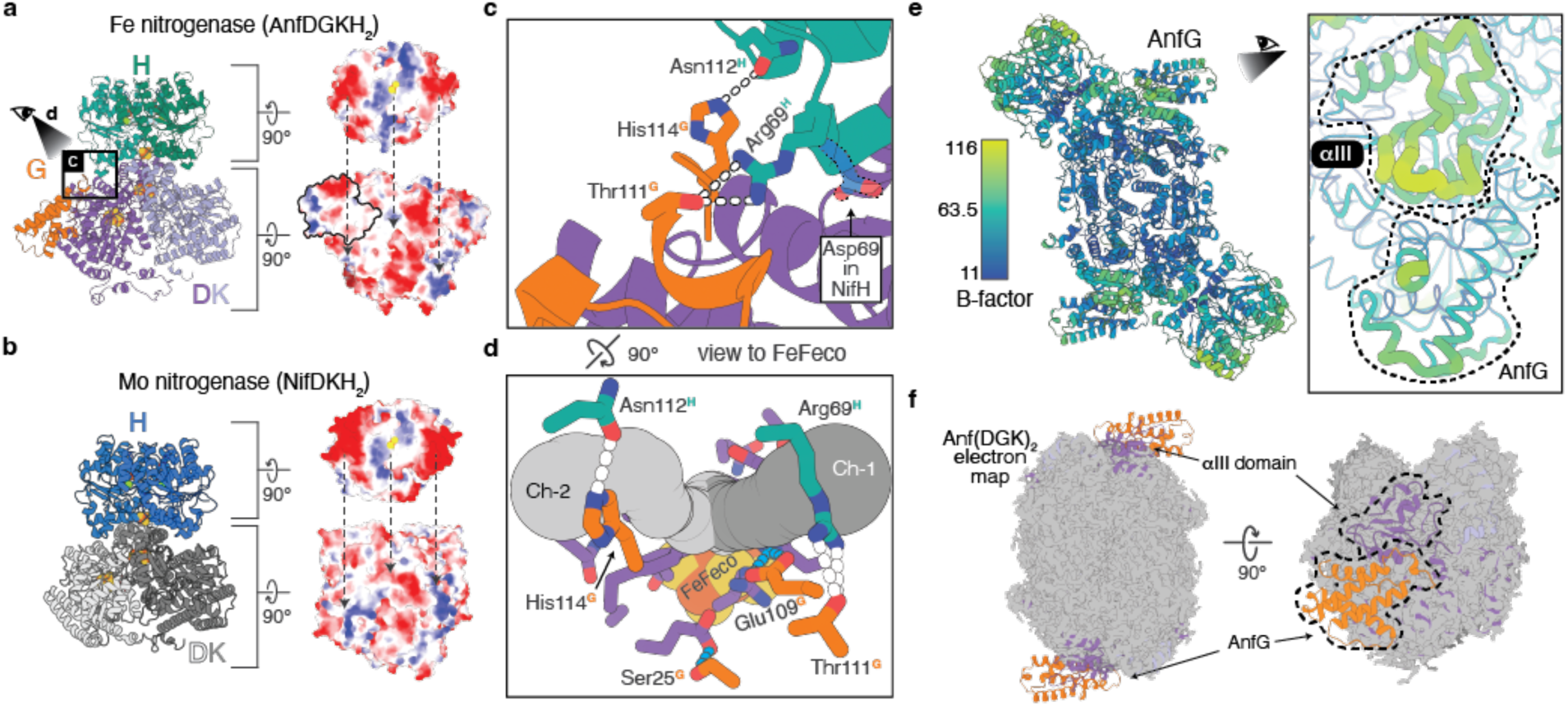
Potential roles of AnfG in the Fe nitrogenase complex. (**a**) Left: Depiction of the AnfDGKH2 subcomplex. Right: Electrostatic potentials of the AnfH2 (top) and AnfDGK (bottom) interaction surfaces. Negative charges are shown in red, neutral in white and positive in blue. Arrows indicate interaction interfaces with complementary charges. AnfG is outlined in black (**b**) Left: Depiction of the NifDKH2 subcomplex (modified from PDB: 7UTA). Right: Electrostatic surface potentials as shown in (a). (**c**) Close-up view on the AnfG – AnfH interaction interface as highlighted in (a). Hydrogen bonds between amino acid residues are indicated by white dashes. Shown in faint blue is Asp69^NifH^ that replaces Arg69^AnfH^ in the Mo nitrogenase complex (PDB: 7UTA), exemplifying the reverse interaction characteristics of the two reductase components. (**d**) 90° rotation from the view in (c) towards the FeFeco showing two CAVER predicted N2 channels to the active site (Ch-1 & -2, tubes in different shades of grey) and a molecular dynamics (MD) calculated N2 channel proposed by Smith et al. [45] (purple residues). Light blue dashes are highlighting interactions of AnfG-residues Ser25^AnfG^ and Glu109^AnfG^ with residues of the MD calculated channel. White dashes depict the same interactions highlighted in (c). (**e**) Left: Per atom B-factors within the Fe nitrogenase complex. Right: Highlight on B-factors in the αIII domain and AnfG in putty representation. (**f**) Model of apo-Anf(DGK)2 fitted into the 2.49 Å cryoEM map of the CHAPSO detergent treated Anf(DGK)2 sample. As highlighted by the arrows, electron density for AnfG and parts of AnfD is missing, including the αIII domain and the FeFeco.

Following complex formation an electron is transferred from the [Fe_4_S_4_]-cluster to the P-cluster, a [Fe_8_S_7_]-cluster embedded at the AnfD-AnfK interface (Fig. 2a and e). In our structure, the [Fe_4_S_4_]-cluster is 17.7 Å apart from the P-cluster. The P-cluster is in the dithionite-reduced P^N^-state [40, 41] forming a symmetric molecule, connected by a shared sulphide ion and bound by six cysteine residues of either the D or K subunit. During catalysis the P-cluster donates electrons to the 19.4 Å distant FeFeco. As previously proposed [13], the FeFeco is a [Fe_8_S_9_C*-(R*)-homocitrate]-cluster (Fig. 2a and f) that in contrast to the Mo and V nitrogenases contains no transition metal other than iron (based on ICP-OES). Six irons (Fe^2^ – Fe^7^) form a trigonal prism around a central carbide that was recently confirmed by X-ray emission spectroscopy [42]. Fe^1^ and Fe^8^ anchor the FeFeco to the AnfD backbone via Cys257^AnfD^ and His423^AnfD^, respectively. The latter iron is additionally coordinated by a bidentate (*R*)-homocitrate ligand that binds the iron atom via its 2-hydroxyl and 2-carboxyl moieties, both with distances of 2.2 Å. Taken together, the FeFeco appears to be almost identical to the FeMoco of the Mo nitrogenase, except for Mo being replaced by another Fe.

In summary, we present the first comprehensive structure of an alternative nitrogenase complex including the reductase and catalytic component. The structure contains two [Fe_4_S_4_]-clusters, two P-clusters and two FeFecos and thus provides direct evidence for the long-hypothesised architecture of the Fe nitrogenase complex.

### Revealing the roles of the G-subunit

The G-subunit is a distinct feature of alternative nitrogenases, but its function remains elusive. Therefore, we analysed our structure for potential roles of the G-subunit in the Fe nitrogenase complex. First, the G-subunit has been proposed to contribute to the specificity between the interaction of the nitrogenase reductase and catalytic component [43]. This interaction is thought to rely mostly on electrostatic interactions [44]. Indeed, when analysing the electrostatic potentials of the AnfH_2_ – Anf(DGK)_2_ interface, we could identify complementary surface charges between the two partner proteins (Fig. 3a). This charge pattern is distinct from the Mo nitrogenase, where the negatively charged regions distal to the [Fe_4_S_4_]-cluster are much more pronounced and the positively charged patch around the reductase cofactor is less accentuated (Fig. 3b). Intriguingly, the G-subunit contributes directly to the binding of AnfH_2_ through hydrogen bonding between His114^AnfG^ and Asn112^AnfH^ as well as Thr111^AnfG^ and Arg69^AnfH^ (Fig. 3c). Instead of the positively charged Arg69^AnfH^, NifH features a negatively charged aspartate residue at the same position, underlining the inverse interaction characteristics of the two reductase components. Hence, AnfG likely determines the specificity between Anf(DGK)_2_ and AnfH_2_ through H-bonding.

Second, AnfG might be involved in directing substrates towards the nitrogenase active site. Recent molecular dynamics (MD) calculations have suggested a N_2_ channel to the Mo nitrogenase active site through the D-subunit [45], which is conserved in the Fe nitrogenase (Fig. 3d). Using the program CAVER [46], we could identify potential substrate channels to the proposed FeFeco substrate binding site at the sulphur S2B [7]. Intriguingly, both the two likeliest CAVER predictions and the MD calculated channel initialise around the AnfG-AnfH interface, the latter even comprising interactions with residues Ser25^AnfG^ and Glu109^AnfG^ (Fig. 3d). These interactions could modulate the channel while leaving enough space for small molecules to enter, thus supporting the idea of a regulatory function of the G-subunit in substrate accessibility.

Third, our data supports an involvement of the G-subunit in stabilising the FeFeco. AnfG is located above the previously described αIII domain, which in the Fe nitrogenase is composed of Arg16^AnfD^ to Lys34^AnfD^ and Tyr359^AnfD^ to Asp384^AnfD^ (Extended Data Fig. 4). The αIII domain forms a lid on top of the active site cofactor that has been shown to undergo major rearrangements during FeMoco insertion [47]. Furthermore, αIII mobility was proposed to play a role in nitrogenase catalysis [27]. Indeed, B-factors around the αIII domain are the highest within the catalytic core of the Fe nitrogenase (Fig. 3e), hinting towards an inherently flexible character of the αIII domain that is stabilised by the interaction with AnfG. To examine if the αIII domain flexibility is observed or even amplified in the resting state of the catalytic component, we tried to solve the cryoEM structure of Anf(DGK)_2_. Using identical conditions as for the AlF_3_-trapped complex, particle orientation had a strong bias for top views on AnfD (Extended Data Fig. 5). Hence, AnfD seems to interact with the air-water interface (AWI) leading to a preferred orientation of the particles. This issue does not occur in our Anf(DGK)_2_(H_2_)_2_ dataset, possibly because the bound reductase component shields the Anf(DGK)_2_-AWI. To circumvent the preferred orientation problem, we collected another dataset of the catalytic component with CHAPSO detergent added right before plunge freezing of the grids. As described previously [48], the use of detergent mitigated the preferred orientation problem, and we obtained a cryoEM map with a global resolution of 2.49 Å (Fig. 3f, Extended Data Fig. 5). However, the map is missing electron density for AnfG, suggesting that it was solubilised from the complex after addition of CHAPSO. Intriguingly, we were not able to resolve electron densities for parts of AnfD in the AnfG-free complex, including the αIII domain and the FeFeco, which is why we did not further refine or deposit this structure. Nevertheless, the G-subunit appears to support FeFeco stabilisation through interactions with the αIII domain and might cover the FeFeco insertion site after cluster insertion.

**Fig. 4:**
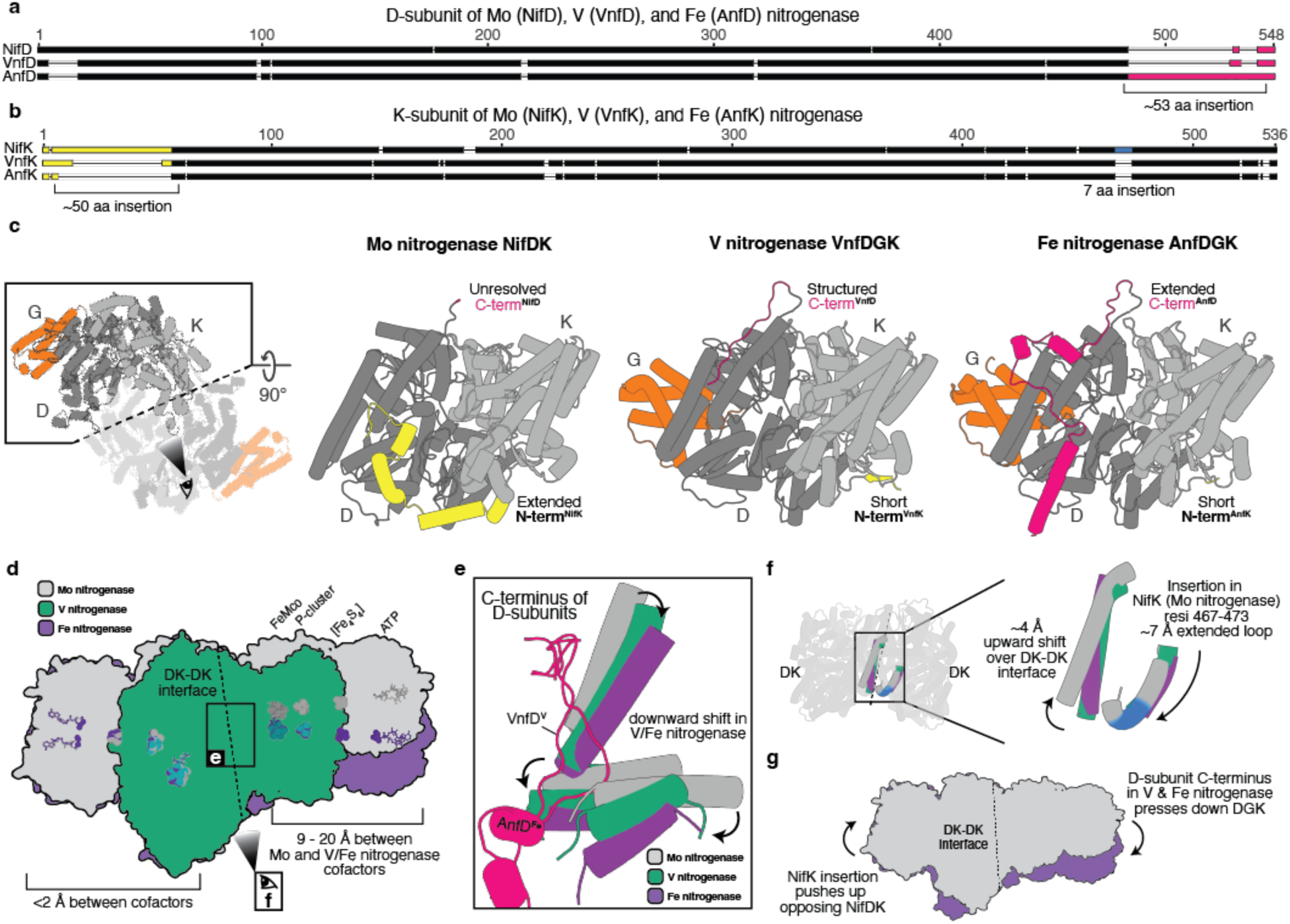
Structural differences between the Mo, V and Fe nitrogenases. (**a**) MUSCLE alignment between NifD (*A. vinelandii*), VnfD (*A. vinelandii*) and AnfD (*R. capsulatus*) amino acid sequences. The C-terminal region is highlighted in magenta, which in AnfD is extended by ∼53 amino acids (**b**) MUSCLE alignment between NifK (*A. vinelandii*), VnfK (*A. vinelandii*) and AnfK (*R. capsulatus*) amino acid sequences. Highlighted in yellow is the N-terminal region, which in NifK is extended by ∼50 amino acids. Highlighted in blue is another seven amino acid insertion in the C-terminal region of NifK. (**c**) View on the DK-DK interface of the nitrogenase catalytic components, including the D and K subunits in different shades of grey and the G-subunits of the alternative nitrogenases in orange. The C-and N-termini are highlighted in magenta and yellow according to (a) and (b). (**d**) Overlay of the Fe nitrogenase (purple) with the Mo nitrogenase (grey, PDB: 7UTA) and the catalytic component of the V nitrogenase (green, PDB: 5N6Y). The respective cofactors are coloured accordingly. The same colour coding applies for (e) - (g) (**e**) Close-up on the DK-DK interface, showing the effects of the respective D-subunit C-termini (highlighted in magenta) on adjacent α-helices. (**f**) View on the DK-DK interface, highlighting the effect of the seven amino acid insertion in the NifK C-terminal region (blue) on the neighbouring α-helix. (**g**) Summary of the effects associated with the observed kink of the Fe nitrogenase relative to the Mo nitrogenase.

Taken together, we propose three roles for the G-subunit in the Fe nitrogenase complex: (*i)* reductase component binding, (*ii)* substrate channelling, and (*iii)* FeFeco insertion and stabilisation.

### Structural comparison of the nitrogenases

Next, we compared the Fe nitrogenase structure to those of the V and Mo nitrogenases. At first glance, the nitrogenase architectures appear to be quite similar with root mean square deviations between individual subunits of less than 3.2 Å for all isoforms and less than 1.4 Å for the two alternative nitrogenases (Extended Data Table 1). However, sequence alignments of the nitrogenase subunits reveal substantial differences in the N-and C-terminal regions of the D and K proteins (Fig. 4a, b). AnfD features an extended C-terminus of approx. 53 amino acids, whereas NifK contains an extended N-terminus of around 50 amino acids and a short seven amino acid insertion in the C-terminal region relative to the other two homologs, respectively. These differences can be observed over a wide range of species, hinting towards a functional relevance of the described features (Extended Data Fig. 6). Intriguingly, all the described features are located at the dimeric interface of DK-DK (Fig. 4b), raising the question whether they could influence the proposed cooperative mechanism between the two halves of the nitrogenase complex [27]. In the Mo nitrogenase, the N-terminal NifK extension wraps around the neighbouring NifD subunit, thereby stabilising the heterodimer (Fig 4c). In contrast, the C-terminal extension of AnfD does not touch the neighbouring AnfK subunit but forms three α−helices that are positioned at the AnfDK-AnfDK interface. Similarly, the VnfD C-terminus is located at the VnfDK-VnfDK interface. However, it is much shorter than the AnfD C-terminus and does not form any secondary structure elements and is more similar to the unstructured C-terminus of NifD [49]. Overlaying our structure with the Mo nitrogenase complex (PDB: 7UTA) and the catalytic component of the V nitrogenase (PDB: 5N6Y) we noticed that the alternative nitrogenases align well with each other, while only one half of the Mo nitrogenase aligns to the Fe and V nitrogenase (Fig. 4d). In the other half, the complexes appear to be kinked relative to each other, with distances between the respective cofactors of up to 20 Å. We could identify two structural differences in the DK-DK interfaces of the three nitrogenases that might cause this effect. On the one hand, the C-terminal regions of the alternative nitrogenases, particularly the extended AnfD C-terminus, wedge themselves into the DK-DK interface (Fig. 4e). Here, they interact with neighbouring α-helices of the respective K-subunits, leading to a downwards shift of the homologous helices in the Mo nitrogenase. On the other hand, the seven amino acid insertion in the NifK C-terminal region (Ile467^NifK^ – Ile473^NifK^) constitutes an extension of the associated α-helix, which pushes an adjacent helix upwards (Fig. 4f). In summary, we propose a complementary effect of the VnfD and AnfD C-termini pressing down and the short NifK insertion pushing up structural elements at the DK-DK interface that cause a kink between the two DK-halves. (Fig. 4g). Thus, the three nitrogenases differ not only in cofactor composition but also show distinct structural features, which may contribute to the unique reactivities observed for the individual isoenzymes.

## Discussion

Here we established *R. capsulatus* as a model organism for the plasmid-based expression and purification of the Fe nitrogenase. After confirming the full N_2_-reduction activity of the purified enzyme, we solved the Fe nitrogenase structure by cryoEM. Due to the oxygen sensitivity of the metalloclusters, preparation of nitrogenase cryoEM samples had to be performed anaerobically, which we successfully accomplished as demonstrated by the reduced P^N^ state of the P-cluster (Fig. 2e). The identified structural features of the Fe nitrogenase complex provide molecular insights into the unique properties of the Fe nitrogenase and highlight general features of the alternative nitrogenases.

One specific feature of the alternative nitrogenases is the presence of an additional α−helical subunit: VnfG and AnfG. Yet, the function of the G-subunit is poorly understood. Based on our structure and additional experiments we suggest three potential roles of AnfG in the Fe nitrogenase complex. (*i)* We identified direct interactions of C-terminal AnfG residues with AnfH_2_, indicating that AnfG is involved in mediating the docking process of the reductase component (Fig. 3c). Previous crystallographic studies have classified three different docking geometries (DG) involved in the electron transfer from the reductase to the catalytic component (DG1 – 3, Extended Data Fig. 7), leading to the hypothesis that the reductase component moves across the surface of the catalytic component during turnover [50, 51]. The ADP-AlF_3_ trapped complex presented here corresponds to the DG2 state, which depicts the moment around the interprotein electron transfer, with AnfH_2_ being in the most central position. In DG3, AnfH_2_ should come even closer to AnfG, which therefore might play a role in energy transduction during nitrogenase catalysis and the release of AnfH_2_ upon ATP hydrolysis. Moreover, the structure indicates that AnfG might contribute to the specificity of AnfH_2_ with Anf(DGK)_2_ (Fig. 3c). This hypothesis is in accordance with previously conducted cross reactivity assays, where N_2_ reduction by the Fe nitrogenase was observed exclusively with AnfH_2_ but not with the two homologous reductase components of *A. vinelandii* [36]. (*ii)* We outlined three potential substrate channels to the FeFeco, which initialise around the location of the G-subunit (Fig. 3d). Therefore, we speculate that AnfG potentially modulates and regulates the substrate access to the active site. This could partially explain the observed reactivity differences of nitrogenase isoforms for N_2_, CO and CO_2_ reduction. (*iii)* Our data suggests that the G-subunit contributes to the FeFeco insertion and stabilisation. This hypothesis is based on our observation that AnfG is located on top of the αIII domain (Fig. 3e), which is associated with the insertion of the active site cofactor and nitrogenase catalysis [27, 47]. We observe that AnfG binds and stabilises the αIII domain, implying that the G-subunit is impacting the processes linked to the αIII domain. In support of this hypothesis, loss of AnfG after the addition of detergent leads to a destabilisation of the αIII domain accompanied by the loss of the FeFeco (Fig. 3f, Extended Data Fig. 5) Thus, the G-subunit might stabilise the active site cofactor through interaction with the αIII domain.

Aligning the Fe with the Mo nitrogenase complex we noticed that the two halves of the catalytic components (NifDK/AnfDGK) are differently interacting with each other leading to a distortion of the catalytic component and a shift of the cofactors in the second half of the nitrogenase complex (Fig 4d – g). In a recent cryoEM study analysing nitrogenase complexes under turnover conditions [27] it has been observed that the two halves of the catalytic component are in different states. Furthermore, only one reductase component was bound at a time, suggesting the catalytic halves to ‘communicate’ with each other to prevent binding of a second reductase component. Due to the divergent interactions of the catalytic halves described here, we expect a different type of “ping-pong” mechanism [27] for the alternative nitrogenases potentially also affecting catalytic rate due to a changed half-reactivity [52]. This could potentially have an influence on the different substrate and product profiles observed for the various nitrogenase isoforms for the reduction of N_2_, CO and CO_2_. A key factor in the communication among the two catalytic halves could be the 53 amino acid extended C-terminus of AnfD that will be the focus of further investigations. In summary, the structure reported herein offers the foundation to rationally modify and test the differences among nitrogenase isoenzymes to provide new insights on the catalytic profiles of the three nitrogenase isoforms.

## Methods

### Chemicals

Unless noted otherwise, all chemicals were purchased from Carl Roth GmbH + Co. KG (Karlsruhe, Germany), Thermo Fisher Scientific Inc. (Waltham, USA), Sigma-Aldrich (St. Louis, USA) or Tokyo Chemical Industry Deutschland GmbH (Eschborn, Germany) and were used directly without further purification. Gases were purchased from Air Liquide (Paris, France).

### Molecular cloning

All used primers were purchased from Eurofins Genomics (Eurofins Scientific SE, Luxembourg City, Luxembourg) and are listed in Table S1. Polymerase chain reactions (PCRs) were conducted with Q5® High-Fidelity DNA Polymerase (New England Biolabs, Ipswich, USA), PCR purifications with the Monarch® PCR & DNA cleanup kit (New England Biolabs, Ipswich, USA), extraction of genomic DNA with the Monarch® Genomic DNA Purification Kit (New England Biolabs, Ipswich, USA), Gibson assemblies with the NEBuilder® HiFi DNA Assembly Master Mix (New England Biolabs, Ipswich, USA) and Golden Gate cloning with the NEBridge® Golden Gate Assembly Kit (New England Biolabs, Ipswich, USA) according to the instructions provided by the manufacturer. Successful assembly of desired vectors was verified by Sanger sequencing through Microsynth Seqlab GmbH (Göttingen, Germany). All plasmids used and created in this study are listed in Table S2.

The pK18mobSacB knockout plasmids were generated via Golden Gate cloning or Gibson assembly. For Gibson assembly, backbone amplification was always done with the primers oMM0227 and oMM0228. The up-and downstream homologous regions of the targeted genomic loci were amplified from *Rhodobacter capsulatus* B10S genomic DNA with primers featuring overhangs suitable for Golden Gate cloning or Gibson assembly. The primers used for the construction of each knockout plasmid are listed in Table S1. The plasmid pBS85-BsaI-*genR*, used for the interruption of the *anfHDGK* locus by a gentamycin resistance cassette, was constructed via Golden Gate cloning. First, a BsaI cutting site was introduced into pBS85 using the primers oMM0027 and oMM0028 to create pBS85-BsaI. Next, Golden Gate inserts were amplified. The gentamycin resistance cassette was amplified from pOGG024 using oMM0033 together with oMM0034. In parallel, the up-and downstream homologous regions of the *anfHDGK* locus were amplified from *R. capsulatus* B10S genomic DNA using oMM0035 – oMM0038. Eventually, the three inserts and the pBS85-BsaI plasmid were combined in a Golden Gate reaction to generate pBS85-BsaI-*genR*.

For the generation of pOGG024-*kanR*, a kanamycin resistance cassette was amplified via PCR from plasmid pRhon5Hi-2 using primers oMM0384 and oMM385. The plasmid pOGG024 was linearized via PCR with oMM0386 and oMM0387. The two DNA amplicons were purified and combined via Gibson assembly, which yielded pOGG024-*kanR*. The construction of the pOGG024-*kanR*-*anfHDGK* expression plasmid was achieved in four steps. First, the *anfHDGK* operon was amplified from *Rhodobacter capsulatus* B10S genomic DNA via PCR using the primers oMM0021 and oMM0146. In parallel, the destination plasmid pRhon5Hi-2 was linearized with oMM0145 and oMM0023. Following purification of the PCR products, pRhon5Hi-2-*anfHDGK* was generated via Gibson assembly. Next, two BsaI cutting sites were removed via a modified version of quick-change mutagenesis [53] using the primers oMM0161 – 164. The resulting plasmid pRhon5Hi-2-*anfHDGK* is suitable for Golden Gate cloning and was used as a template to amplify the *anfHDGK* expression cassette with the primers oMM0389 and oMM0390. The resulting PCR product was purified and subsequently inserted into pOGG024-*kanR* via Golden Gate cloning. Lastly, affinity tags (Strep-tag II and (His)_6_-tag) for protein purification were inserted via restriction free cloning [54] using primers oMM0223 plus oMM0224 for the Strep-tag II and oMM0510 plus oMM0511 for the (His)_6_-tag. Eventually, the sequence of the pOGG024-*kanR*-*anfHDGK* expression plasmid was confirmed by whole plasmid sequencing through plasmidsaurus (Eugene, USA).

### Genetic manipulation of *Rhodobacter capsulatus*

Starting from the wildtype strain B10S, the *Rhodobacter capsulatus* (*R. capsulatus*) genome was successively modified to generate an ideal strain for the recombinant expression and subsequent purification of the Fe nitrogenase. For the deletion of *anfHDGK*, a gentamycin resistance cassette was inserted into the *anfHDGK* locus, thereby interrupting the operon. The plasmid pBS85-BsaI-*genR* was introduced into *R. capsulatus* B10S via conjugational transfer as described in [29], selecting for the gentamycin resistance conferred by the transferred vector. Subsequently, obtained clones were screened for gentamycin resistance and tetracycline sensitivity on peptone yeast (PY) agar plates [29] containing 15 µg/mL gentamycin or 10 µg/mL tetracycline, respectively. Positive clones were further investigated via colony PCR to check the *anfHDGK* locus. The purified PCR products were analysed by Sanger Sequencing to identify clones with a successfully modified *anfHDGK* operon. Building up on the Δ*anfHDGK::genR* mutant of *R. capsulatus* B10S, all further deletions were achieved successively via the *sacB* scar less deletion method described in [55]. In brief, sequences of around 500 base pairs homologous to the up-and downstream regions flanking the gene of interest (GOI) were generated and cloned into a pK18mobSacB suicide vector (see above). The resulting plasmid was conjugated into the *R. capsulatus* recipient strain [29], selecting for the kanamycin resistance conferred by the suicide vector. Intermediate strains derived from single colonies that were obtained from the previous step were passaged three times in liquid peptone yeast (PY) medium [29], growing each passage for 24 h at 30°C and moderate shaking in the dark. The final passage was spread on a PY agar plate containing 5% (m/V) sucrose, which was then incubated for 72 h at 30°C under an argon atmosphere and illumination by six 60 W krypton lamps (Osram Licht AG, Munich, Germany). Single colonies of *R. capsulatus* growing on the sucrose containing agar plate were screened for kanamycin and sucrose sensitivity on PY plates containing 50 µg/mL kanamycin or 5% (m/V) sucrose, respectively. Colonies that could tolerate sucrose but were not growing on kanamycin containing agar plates were further investigated via colony PCR to check the targeted genomic locus. Lastly, the purified PCR products were analysed by Sanger sequencing (Microsynth Seqlab GmbH, Göttingen, Germany) to identify successful knockout clones. Genomic DNA of the modified *R. capsulatus* B10S strain (MM0425) was extracted and sequenced via next generation sequencing (Novogene Co., Ltd., Beijing, China) to confirm the deletions listed in Table 1. The *R. capsulatus* MM0436 expression strain was generated by introducing the pOGG024-*kanR*-*anfHDGK* expression plasmid into MM0425 via conjugational transfer, all used strains are listed in Table S3.

**Table 1:** Genetic modifications of *Rhodobacter capsulatus* expression strain.

| Modification | Deleted locus naturally encodes for | Purpose for deletion | Ref. |
| --- | --- | --- | --- |
| $\Delta nifHDK$ | Molybdenum nitrogenase | Remove the primary nitrogenase | [34] |
| $\Delta modABC$ | High affinity molybdenum transporter | Prevent molybdenum import to maximise expression levels of the alternative nitrogenase | [31] |
| $\Delta draTG$ | ADP-ribosyltransferase/ADP-ribosyl hydrolase system for the post translational modification of nitrogenase iron proteins | Ensure constitutive activity of the Fe-only nitrogenase | [32] |
| $\Delta gtaI$ | Quorum sensing protein GtaI | Removes the capsule to increase cell pellet quality post centrifugation | [33] |
| $\Delta anfHDGK::gmR$ | Fe-only nitrogenase | Prevent genomic expression of wild-type Fe-only nitrogenase to enable recombinant expression of AnfHDGK variants from plasmid DNA | [56] |

### Growth medium and conditions for protein production

*R. capsulatus* was cultivated phototrophically at 32 °C under a 100% dinitrogen (N_2_) atmosphere. Cultivation on agar plates was conducted on peptone yeast (PY) agar plates [29] selective for the respective expression plasmid. Liquid cultures of *R. capsulatus* were cultivated diazotrophically in a modified version of RCV medium [29] that contained 30 mM DL-malic acid, 0.8 mM MgSO_4_, 0.7 mM CaCl_2_, 0.05 mM sodium ethylenediaminetetraacetic acid (Na_2_EDTA), 0.03 mM thiamine hydrochloric acid, 9.4 mM K_2_HPO_4_, 11.6 mM KH_2_PO_4_, 5 mM serine, 1 mM Fe(III)-citrate, 45 µM B(OH)_3_, 9.5 µM MnSO_4_, 0.85 µM ZnSO_4_, 0.15 µM Cu(NO_3_)_2_ and 25 µg/mL kanamycin sulphate at a pH set to 6.8. For protein production, the expression strain was inoculated from a glycerol stock on PY agar plates and cultivated for 48 h. Obtained cell mass was used to inoculate liquid cultures in N_2_-flushed RCV medium, which were cultivated for 24 h. Subsequently, the cultures were diluted into 800 mL RCV medium to an optical density of 0.1 at 660 nm (OD_660_). Protein purification was initiated when the cultures reached an OD_660_ of ∼ 3.0.

### Protein purification

All protein purification steps were carried out strictly anaerobically under a 95% argon (Ar) 5% dihydrogen (H_2_) atmosphere inside a COY tent (Coy Laboratory Products, Inc. Grass Lake, USA). All buffers were anaerobised by flushing them with Ar and equilibrating them for at least 12 h inside the COY tent before use. For harvesting, sodium dithionite was added to a final concentration of 5 mM to each liquid culture, which were then centrifuged at 15970 × g for 60 min, 10°C. The liquid supernatant was decanted and the cell pellets were resuspended and combined in high salt buffer (50 mM TRIS (pH = 7.8), 500 mM NaCl, 10% glycerol, 4 mM sodium dithionite) supplemented with 0.2 mg/mL bovine pancreatic deoxyribonuclease I and one cOmplete EDTA-free protease inhibitor tablet (Roche, Basel, Switzerland). Subsequently, cells were disrupted by three passages through a French press cell disrupter (catalogue #FA-078AE; Thermo Fisher Scientific Inc., Waltham, USA) at 20000 psi. The obtained lysate was centrifuged for 60 min at 150,000 × g and 8°C and the liquid supernatant was filtered (pore size = 0.2 µm). The cleared cell extract was then applied to high salt buffer equilibrated HisTrap™ HP (Cytiva Europe GmbH, Freiburg, Germany) and Strep-Tactin®XT 4Flow® high capacity (IBA Lifesciences, Göttingen, Germany) columns via a ӒKTA pure™ chromatography system (Cytiva Europe GmbH, Freiburg, Germany). After extensive washing with binding buffer, the catalytic component was eluted from the Strep-Tactin®XT column with binding buffer supplemented with 50 mM biotin. Fractions containing the catalytic component were pooled, buffer exchanged to low salt buffer (50 mM TRIS (pH = 7.8), 150 mM NaCl, 10% glycerol, 4mM sodium dithionite) with a Sephadex G-25 packed PD-10 desalting column (Cytiva Europe GmbH, Freiburg, Germany) and concentrated with an Amicon® Ultra-15 Centrifugal Filter Unit (molecular weight cut off = 100 kDa; Merck Millipore, Billerica, USA). Meanwhile, the HisTrap™ HP column was washed extensively with high salt buffer containing 25 mM imidazole before eluting the reductase component with high salt buffer plus 250 mM imidazole. The eluate was directly subjected to a size exclusion chromatography on a HiLoad 26/600 Superdex 200 pg column (Cytiva Europe GmbH, Freiburg, Germany) equilibrated with low salt buffer. AnfH_2_ eluted in a clear peak at around 205 mL and was subsequently concentrated using an Amicon® Ultra-15 Centrifugal Filter Unit (molecular weight cut off = 30 kDa; Merck Millipore, Billerica, USA). Protein yields for both nitrogenase component fractions were determined using the Quick Start™ Bradford 1x Dye Reagent (Bio-Rad Laboratories, Inc., Hercules, USA) according to the instructions by the manufacturer and purity of both protein fractions was analysed via sodium dodecyl sulphate polyacrylamide gel electrophoresis (SDS-PAGE). Eventually, both nitrogenase components were flash frozen and stored in liquid N_2_ until further use.

### SDS-PAGE analysis

For sodium dodecyl sulphate polyacrylamide gel electrophoresis (SDS-PAGE), protein samples were denatured by boiling them with Pierce™ Lane Marker Reducing Sample Buffer (Thermo Fisher Scientific Inc., Waltham, USA) at 98°C for 10 min. After centrifuging the samples at 17,000 × g, the clear supernatant was loaded on a 4–20% Mini-PROTEAN TGX Stain-Free Gel (Bio-Rad Laboratories, Inc., Hercules, USA) including PageRuler™ Plus Prestained Protein Ladder (Thermo Fisher Scientific Inc., Waltham, USA) as a molecular weight reference. The electrophoresis was run at a constant voltage of 180 V for 30 min before staining the gel with GelCode™ Blue Safe Protein Stain (Thermo Fisher Scientific Inc., Waltham, USA).

### Nitrogenase turnover assays

Nitrogenase activity was assessed *in vitro* by measuring specific activities for dihydrogen (H_2_) and ammonia (NH_3_) formation under a dinitrogen (N_2_) or argon (Ar) atmosphere. Working under an Ar atmosphere, varying amounts of AnfH_2_ were dissolved in an anaerobic solution of 50 mM TRIS (pH = 7.8), 10 mM sodium dithionite, 3.5 mM adenosine triphosphate (ATP), 7.87 mM MgCl_2_, 44.59 mM creatine phosphate and 0.20 mg/mL creatine phosphokinase (catalogue #C3755; Sigma-Aldrich St. Louis, USA). The reaction vials were sealed by crimping them with butyl rubber stoppers and the headspace was exchanged to N_2_ or Ar. Next, the reactions were initialised by adding 0.1 mg Anf(DGK)_2_ up to a total volume of 700 µL and allowed to proceed for 8 min at 30°C and moderate shaking at 250 rpm. Reactions were quenched with 300 µL 400 mM sodium ethylenediaminetetraacetic acid solution (pH = 8.0) and the amounts of formed H_2_ and NH_3_ were analysed as described below.

### Quantification of dihydrogen

Amounts of formed dihydrogen (H_2_) were determined via headspace analysis using a Clarus®690 GC system (GC–FID/TCD; PerkinElmer Inc., Waltham, USA) with a custom-made column circuit (ARNL6743). The headspace samples were injected by a TurboMatrixX110 (PerkinElmer Inc., Waltham, USA) auto sampler, heating the samples to 45 °C for 15 min prior to injection. The samples were then separated on a HayeSep column (7’ HayeSep N 1/8’’ Sf; PerkinElmer Inc., Waltham, USA), followed by molecular sieve (9’ Molecular Sieve 13x 1/8’’ Sf; PerkinElmer Inc., Waltham, USA) kept at 60 °C. Subsequently, the gases were detected with a flame ionization detector (FID, at 250 °C) and a thermal conductivity detector (TCD, at 200 °C). The quantification of H_2_ was based on a linear standard curve that was derived from measuring varying amounts of H_2_ under identical conditions.

### Quantification of ammonia

Quantification of *in vitro* generated ammonia (NH_3_) was done with a modified version of a fluorescence NH_3_ quantification method described in [57]. 100 µL sample were combined with 1 mL of a solution containing 2 mM o-phthalaldehyde, 10 % (V/V) ethanol, 0.05 % (V/V) β-mercaptoethanol and 0.18 M potassium phosphate buffer (pH = 7.3) and incubated at 25°C for 2 h in the dark. 50 µL of each sample were transferred into individual wells of a black Nunc™ F96 MicroWell™ plate (Thermo Fisher Scientific Inc., Waltham, USA) and fluorescence at 485 nm was monitored with an Infinite® 200 PRO plate reader (Tecan Group Ltd, Männedorf, Switzerland) in fluorescence top reading mode using an excitation wavelength of 405 nm. The quantification of ammonia was based on a linear standard curve that was derived from measuring varying amounts of NH_4_Cl under identical conditions. Samples incubated under an argon atmosphere instead of dinitrogen were used to correct for background signal.

### Mass photometry

Mass photometry measurements were carried out on microscope coverslips (1.5 H, 24 × 50 mm; Carl Roth GmbH + Co. KG, Karlsruhe, Germany) with CultureWell^TM^ Reusable Gaskets (CW-50R-1.0, 50-3mm diameter × 1 mm depth) that had been washed with three consecutive rinsing steps of distilled H_2_O and 100% isopropanol and dried under a stream of pressurized air. Measurements were set up in gaskets assembled on microscope coverslips on the stage of a TwoMP mass photometer (MP, Refeyn Ltd, Oxford, UK) with immersion oil. Samples were measured in anaerobic measurement buffer (150 mM NaCl, 50 mM TRIS (pH = 7.8), 10% Glycerol, 10 mM sodium dithionite) after focusing on the glass surface using the droplet-dilution focusing method. After focusing, 0.5 µL nitrogenase sample (500 nM stock concentration, dissolved in measurement buffer with 4 mM dithionite) was removed from an anaerobic vial, quickly added to 19.5 µL measurement buffer, and mixed on the stage of the MP. Measurements were started ∼5 s after removing protein from the anaerobic environment. Data was acquired for 60 s at 100 frames per second using AcquireMP (Refeyn Ltd, Oxford, UK). MP contrast was calibrated to molecular masses using 50 nM of an in-house purified protein mixture containing complexes of known molecular mass. MP datasets were processed and analyzed using DiscoverMP (Refeyn Ltd, Oxford, UK). The details of MP image analysis have been described previously [58].

### Metal analysis

Metal analysis was done via inductively coupled plasma optical emission spectroscopy (ICP-OES). For sample preparation, 0.12 mg and 0.24 mg of catalytic and reductase component, respectively, were dissolved in 0.5 mL trace metal grade concentrated nitric acid and incubated for 12 h at 25°C. Subsequently, the samples were boiled at 90°C for 2 h before they were diluted 17-fold in distilled water. The metal content was analysed with a 720/725 ICP OES device (Agilent Technologies Inc., Santa Clara, USA) on iron (λ = 238.204 nm), molybdenum λ = 202.032 nm), nickel (λ = 216.555 nm) and zinc (λ = 213.857 nm). All analysed metals were quantified using ICP multi-element standard solution IV (Merck KGaA, Darmstadt, Germany) as a standard.

### Preparation of aluminium fluoride stabilised nitrogenase complex

Stabilised Fe nitrogenase complex consisting of two reductase and one catalytic component was prepared as described in [59]. In brief, 4 nmol catalytic and 32 nmol reductase component were combined in 100 mM MOPS, 50 mM TRIS, 100 mM NaCl (pH = 7.3) with 5 mM sodium dithionite, 4 mM NaF, 0.2 mM AlCl_3_, 8 mM MgCl_2_ and 1 mM ATP in a total volume of 4 mL. The reactions were incubated at 30°C for 1 h before they were concentrated with an Amicon® Ultra-0.5ml Centrifugal Filter Unit (molecular weight cut off = 100 kDa; Merck Millipore, Billerica, USA). Subsequently, less than 500 µL sample were injected via a ӒKTA pure™ chromatography system (Cytiva Europe GmbH, Freiburg, Germany) onto a Superdex 30 Increase 10/300 GL column (Cytiva Europe GmbH, Freiburg, Germany) equilibrated with 50 mM TRIS (pH = 7.8), 200 mM NaCl and 5 mM sodium dithionite. Elution fractions from the peak corresponding to the appropriate molecular weight species (expected molecular weight of catalytic component combined with two reductase components is around 372 kDa) were pooled and the presence of all nitrogenase subunits was confirmed via SDS-PAGE as described above.

### CryoEM sample preparation and data collection

4 µL of protein solution (total protein concentration = 1 mg/mL) were applied to freshly glow-discharged QUANTIFOIL® R2/1 300 mesh grids (Quantifoil Micro Tools GmbH, Großlöbichau, Germany) and blotted for 5 s with a blot force of 5 at ∼90% humidity and 8°C using a Vitrobot Mark IV (Thermo Fisher Scientific Inc., Waltham, USA) that was placed inside an anaerobic COY tent. In case of CHAPSO detergent supplemented grids, 1 µL of detergent (dissolved in the same buffer as the protein) was added to a final concentration of 0.4% (m/V) to 3 µL protein solution on the respective. Grids were plunge frozen in a liquid ethane (37 vol%) propane (63 vol%) mix and stored in liquid nitrogen until data collection. CHAPSO supplemented grids of AnfDGK were prepared to prevent preferential orientation.

Data of cryoEM samples were collected on a Titan Krios G3i electron microscope (Thermo Scientific), at an acceleration voltage of 300 kV and equipped with a BioQuantum energy filter (Gatan) and a K3 direct electron detector (Gatan). Data were collected in electron counting super-resolution mode at a nominal magnification of 105,000x (0.837 Å per pixel) with a total dose of 50 e−/Å^2^ (50 fractions), using the aberration-free image-shift (AFIS) correction in the EPU software (Thermo Scientific). The nominal defocus range used for data collection was -1.4 to -2.4 μm.

### CryoEM data processing

All datasets were processed entirely in CryoSparc [60]. For all datasets dose-fractionated movies were gain-normalized, aligned, and dose-weighted using Patch Motion correction and the contrast transfer function (CTF) was determined using the Patch CTF routine. The information regarding CryoEM data collection model refinement and statistics are listed in Table S4.

#### Processing the AnfHDGK-AlF_3_ Complex

Blob picker and manual inspection of particles were used to extract an initial 2,114,475 particles with a box size of 300 pixels, which were used to build 2D classes. 2D classes with protein-like features were used to initialize template picking. After manual inspection and extraction with a box size of 300 pixels, this yielded a total of 3,365,366 particles, which were used to build 2D classes. After selecting 2D classes with protein-like features, the selected particles were used to train a model that was subsequently used to pick particles using TOPAZ [61]. A total of 1,706,699 candidate particles were extracted with a box size of 380 pixels and cleaned from non-particle candidates by 2D classification into 200 classes. Selected particles were used for *ab-initio* reconstruction and classification into 4 classes. Particles of the 2 best aligning classes (432,216 particles) were subjected to further cleaning by 3D classification into 10 classes with a target resolution of 5 Å. 3D classification yielded volumes containing 0, 1, or 2 AnfG subunits, with unchanged orientation of the remaining subunits. The best aligning classes with 1 or more AnfG subunit bound (218,653 particles) were subjected to local CTF refinement, local motion correction, and subsequent non-uniform refinement with C2 symmetry, 2 extra final passes, 15 Å initial lowpass resolution, 12 Å GSFSC split resolution, 4 Å dynamic mask near expansion, 10 Å dynamic mask far expansion, 8 Å dynamic mask start resolution, per-particle defocus optimization, and EWS correction, yielding 2.35 Å global resolution and a temperature factor of -76.7 Å^2^. Further classification did not yield improved resolution.

#### Processing the AnfDGK component

Initial attempts to solve the AnfDGK complex structure without reductase component used grids prepared without detergent (CHAPSO). Standard processing workflows of this dataset (blob picking, template picking, TOPAZ picking and manual picking) yielded 2D classes that exclusively showed one orientation (Extended Data Figure 5a). Resulting *ab-initio* and 3D-reconstructions failed to yield initial volumes with a nitrogenase-like shape. We therefore focused our efforts on grids prepared in presence of 0.5% CHAPSO.

Here, blob picker and manual inspection of particles were used to extract an initial 2,018,560 particles with a box size of 320 pixels from 2000 micrographs, which were used to build 2D classes. 2D classes with protein-like features were used to train a TOPAZ model to pick particles, which was subsequently used to re-extract particles from the same 2000 micrographs for downstream 2D classification and TOPAZ model training. A total of 1,647,264 particles were extracted with a box size of 340 pixles and cleaned from non-particle candidates by 2D classification. Cleaned particles were used to train a TOPAZ model on 4578 micrographs and subsequently used to pick particles from all 18320 micrographs. A total of 7.962,489 particles were extracted with a box size of 324 pixels and cleaned by three subsequent rounds of 2D classification into 200, 100 and 50 classes, respectively (Extended Data Figure 5b). Selected particles of the last 2D classification step (2,121,950) were used for *ab-initio* reconstruction and classification into 4 classes. Particles of the 2 best-aligning classes (1,336,362 particles) were subjected to further cleaning by 3D classification into 10 3D classes with a target resolution of 4 Å. 3D classification yielded no volumes containing electron density at positions where AnfG would be expected. Nevertheless, particles of the three best-aligning classes (304,619 particles) were used for non-uniform refinement with C1 symmetry and no additional corrections. This yielded a 2.64 Å global resolution map that contained no indication of electron density at locations where AnfG would be expected, nor at select regions of AnfDK in close contact with AnfDK. A subsequent non-uniform refinement using particles of the 7 best-aligning classes (563,245) from the 3D classification, the 2.64 Å map as an input volume, C2 symmetry, CTF-, defocus-and Ewald sphere correction yielded a map with a global resolution of 2.49 Å. This map also contained no electron density at locations where AnfG would be expected, nor at regions in AnfDK that would be near the expected AnfG position. Further classification was not attempted given that AnfG could not be detected in processed volumes.

### Model building and refinement

Initial cryoEM map fitting was performed in UCSF-Chimera 1.16 [62] using AlphaFold [35] models for AnfD, AnfK, and AnfG, as well as an AnfH crystal structure (PDB: 7QQA) from *Azotobacter vinelandii* [36]. The resulting model was manually built further in COOT 0.8.9.2 [63]. Automatic refinement of the structure was done using phenix.real_space_refine of the PHENIX 1.21.1 software suite [64]. Manual refinements as well as water picking were performed with COOT. The FeFeco was built with REEL of the PHENIX software suite. The model statistics are listed in Table S4.

### Substrate Channel Calculation

Substrate cannels were calculated using the software CAVER [46]. The coordinates of sulphur atom S2B were provided as the starting point for channel calculations. The probe radius, shell radius, and shell depth were set to 0.7, 4.0, and 5.0 Å, respectively. Many channels were predicted by CAVER. However, the two most probable channels with the shortest length, the largest bottleneck radius, the highest throughput, prioritized by CAVER were selected and are displayed throughout the manuscript as surfaces generated in PyMOL (Fig. 3d).

## Material availability

All unique materials used in this study are available from the corresponding author upon request.

## Data availability

All raw data for mass photometry measurements, kinetic experiments, and protein characterisation will be deposited on Edmon, the Open Research Data Repository of the Max Planck Society for public access. The atomic structure reported in this paper is deposited to the Protein Data Bank under accession code 8OIE. CryoEM data were deposited to the Electron Microscopy Data Bank under EMD-16890.

## Acknowledgements

The authors thank the Central Electron Microscopy Facility at the Max Planck Institute of Biophysics for expertise and access to their instruments. We thank S. Freibert and J.M. Schuller for help with anaerobic plunge freezing of cryo-EM sample grids and the use of their equipment. We thank C. Thölken, P. Klemm and M. Lechner for help with data management and computing cluster access. We thank S. Reinhard for aid in data and sample transport. We thank M. Girbig, F. Ramirez, A. Kumar and J.M. Schuller for help during cryoEM data processing.

## Author information

These authors contributed equally: Frederik V. Schmidt, Luca Schulz

### Authors and Affiliations

*Research Group Microbial Metalloenzymes, Max-Planck-Institute for Terrestrial Microbiology; Karl-von-Frisch Straße 10, 35043 Marburg, Germany*

Frederik V. Schmidt, Niels N. Oehlmann, Johannes G. Rebelein

*Department of Biochemistry & Synthetic Metabolism, Max-Planck-Institute for Terrestrial Microbiology; Karl-von-Frisch Straße 10, 35043 Marburg, Germany*

Luca Schulz, Jan Zarzyck, Tobias J. Erb

*Central Electron Microscopy Facility, Max-Planck-Institute for Biophysics; Max-von-Laue-Straße 3, 60438 Frankfurt am Main, Germany* Simone Prinz

### Contributions

J.G.R. conceived and supervised the project. T.J.E. and J.G.R. acquired funding. F.V.S., L.S. and J.G.R. designed and analysed experiments. F.V.S. and N.N.O. performed molecular work. F.V.S. performed anaerobic protein purification and enzyme biochemistry. F.V.S. and L.S. performed mass photometry experiments. F.V.S., L.S. and S.P. performed cryoEM data acquisition. L.S., F.V.S. and J.Z. processed and refined the cryoEM structure. L.S., F.V.S., J.Z. and J.G.R. analysed the cryoEM structure. F.V.S., L.S. and J.G.R. wrote the original manuscript which was reviewed and edited by all authors.

### Corresponding author

Correspondence to Johannes G. Rebelein

## Extended Data

**Extended Data Table 1:** Root-mean square deviation between subunits from different nitrogenases. *Azotobacter vinelandii* Mo and V nitrogenase (PDB ID 7UTA and 5N6Y, respectively) were aligned to each other and to *Rhodobacter capsulatus* Fe nitrogenase (PDB: 8OIE).

| D | Mo | V | Fe | K | Mo | V | Fe | G | V | Fe |
| --- | --- | --- | --- | --- | --- | --- | --- | --- | --- | --- |
| Mo | X | 2.72 | 2.67 | Mo | X | 3.11 | 2.84 | V | X | 1.35 |
| V | 2.72 | X | 1.04 | V | 3.11 | X | 1.23 | Fe | 1.35 | X |
| Fe | 2.67 | 1.04 | X | Fe | 2.84 | 1.23 | X |  |  |  |

**Extended Data Fig. 1:**
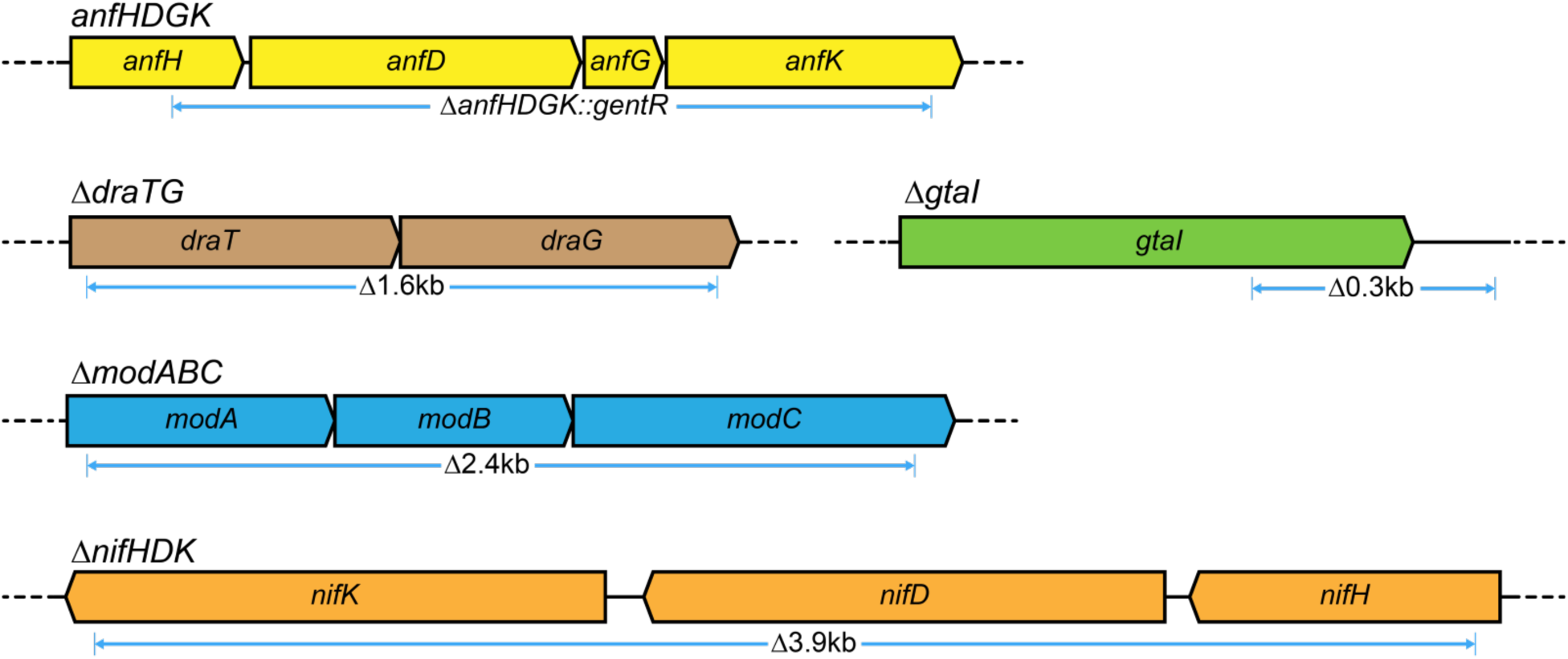
Modified genomic regions and the corresponding polymorphism derived NGS data. Strain MM0425 was sequenced using Illumina sequencing (Novogene Co., Ltd., Beijing, China). The individual reads were trimmed, paired and assembled to the *R. capsulatus* reference genome (Strain SB1003, GenBank CP001312.1) using the Breseq pipeline.

**Extended Data Fig. 2:**
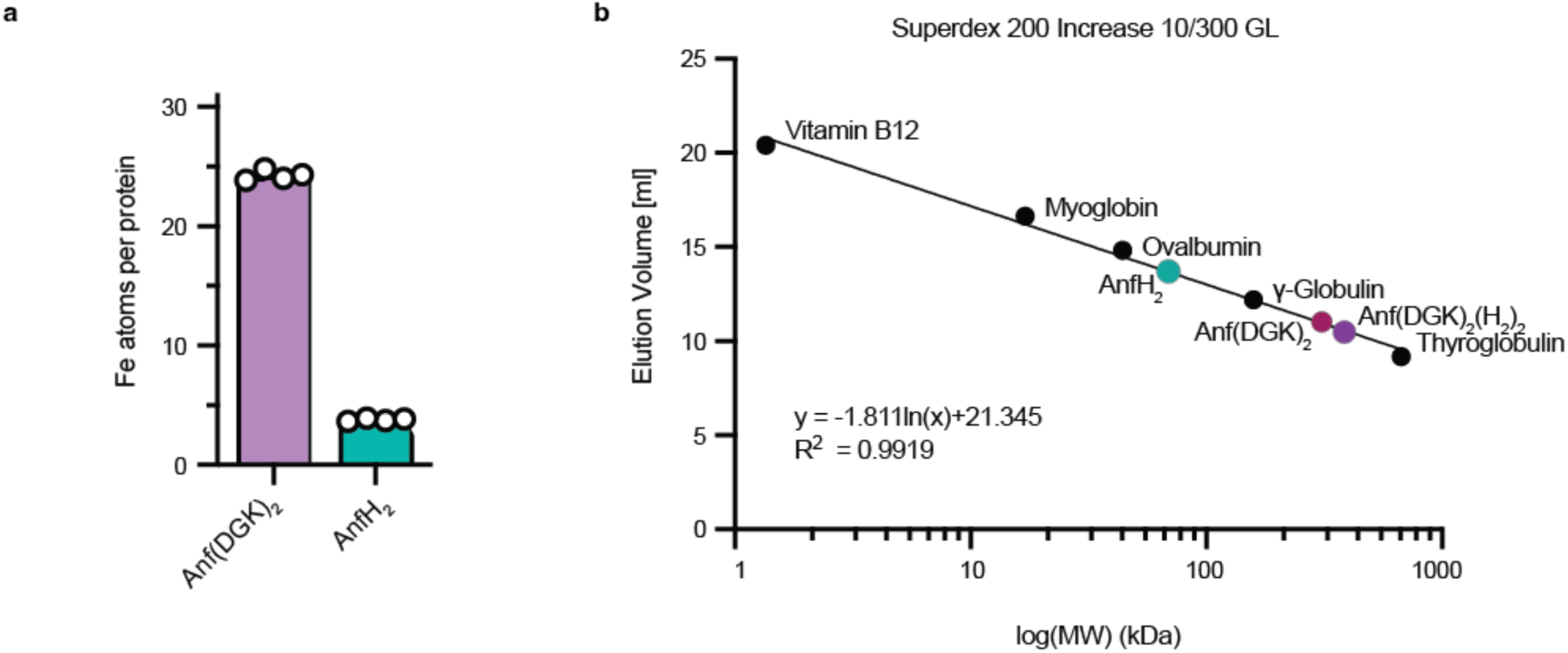
Analysis of Anf(DGK)_2_ and AnfH_2_. (**a**) Inductively coupled plasma optical emission spectroscopy (ICP-OES) data for Anf(DGK)_2_ and AnfH_2_. Data are 2 technical replicates of 2 biological replicates and error bars represent standard deviation. (**b**) Analytical size-exclusion chromatography standard run on a Superdex 200 Increase 10/300 GL column (Cytiva Europe GmbH, Freiburg, Germany) used to infer the complex masses of AnfH_2_, Anf(DGK)_2_ and Anf(DGK)_2_(H_2_)_2_. Black dots indicate proteins included in the standard mixture; coloured dots indicate measured complexes.

**Extended Data Fig. 3:**
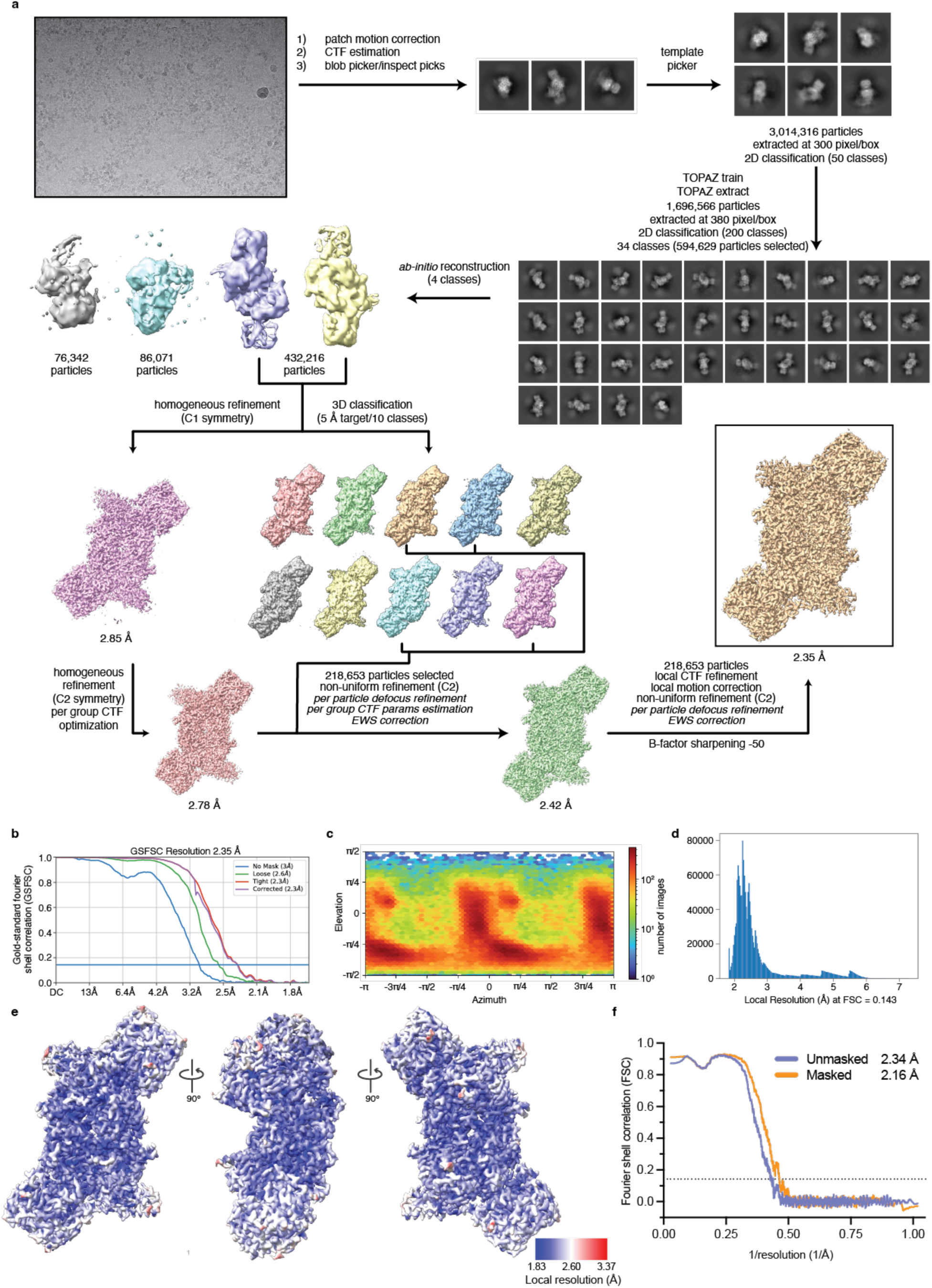
Cryogenic electron microscopy data collection and analysis of Anf(DGK)_2_(H_2_)_2_. (**a**) Schematic data processing workflow for the electron map of Anf(DGK)_2_(H_2_)_2_. Dataset was collected on a Titan Krios G3i electron microscope operated at an acceleration voltage of 300 kV and equipped with a BioQuantum energy filter and a K3 direct electron detector. Dataset was processed entirely in CryoSPARC [60]. (**b**) Gold-standard Fourier shell correlation plot from map refinement in CryoSPARC. Resolution determined at Fourier shell correlation (FSC) = 0.143. (**c**) Angular particle distribution. (**d**) Distribution of local resolution at FSC = 0.143. (**e**) Local resolution as calculated by CryoSPARC mapped onto the refined density with different views shown. (**f**) Map to atomic model FSC plot determined at FSC = 0.143.

**Extended Data Fig. 4:**
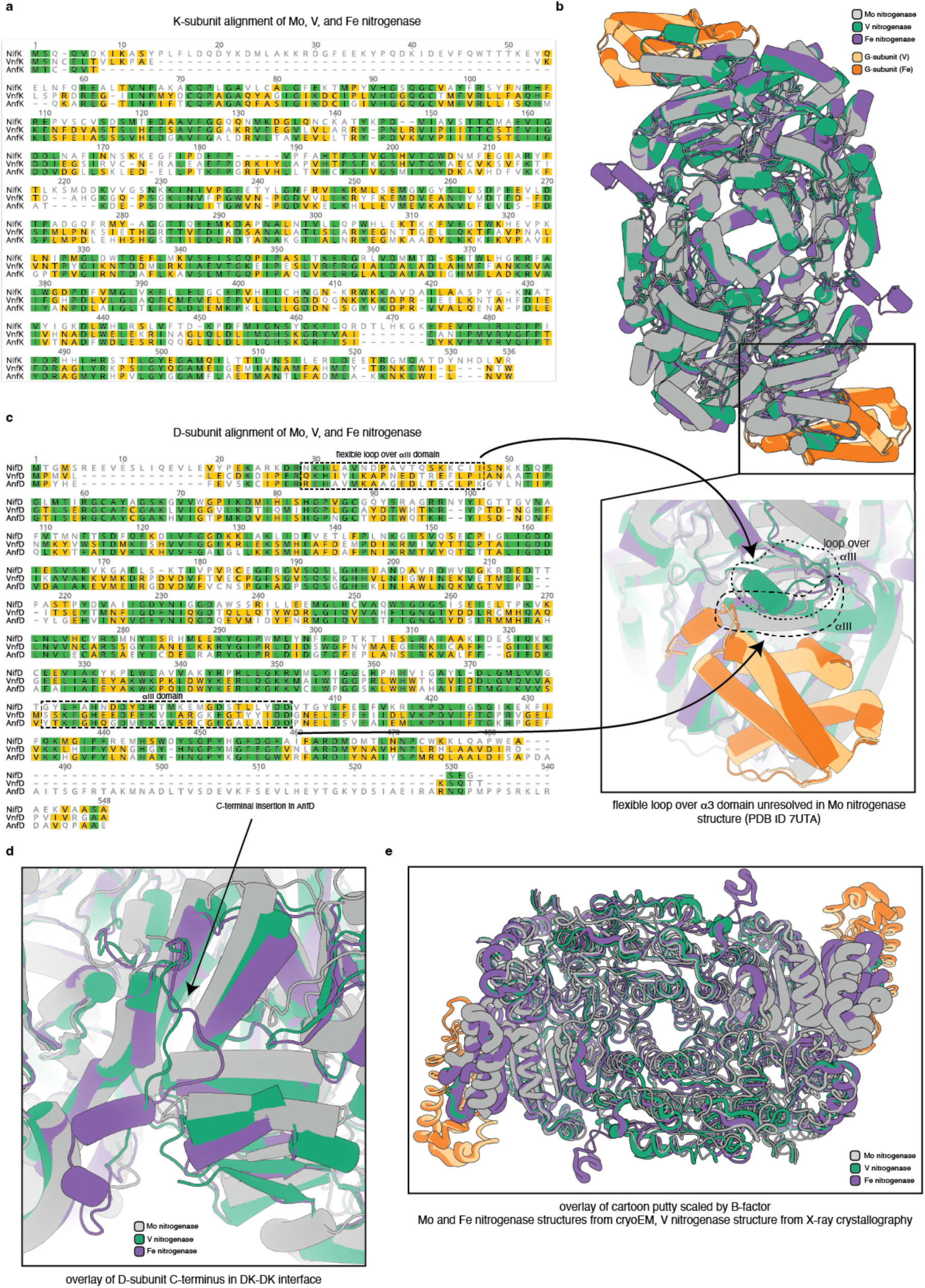
Sequence and structural alignment of Mo, V, and Fe nitrogenase catalytic components. (**a**) Sequence alignment of *Azotobacter vinelandii* Mo and V nitrogenase K-subunit (NifK and VnfK) to *Rhodobacter capsulatus* Fe nitrogenase K-subunit (AnfK). Identical sites are shown in green and similar sites (BLOSUM62 matrix [65] threshold of 2) that occur in 2 of 3 sequences are highlighted in yellow. (**b**) Structural alignment of the catalytic components of the Mo (PDB: 7UTA), V (PDB: 5N6Y), and Fe (PDB: 8OIE) nitrogenases aligned in (a) and including the G-subunit for V and Fe nitrogenase. Inset at bottom shows a zoomed view of the a-III domain, with arrows depicting the origin in the sequence alignment. (**c**) Sequence alignment of *Azotobacter vinelandii* Mo and V nitrogenase D-subunit (NifD and VnfD) to *Rhodobacter capsulatus* Fe nitrogenase D-subunit (AnfD). Identical sites are shown in green and similar sites (BLOSUM62 matrix, threshold of 2) that occur in 2 of 3 sequences are highlighted in yellow. (**d**) Close-up view into the interface between individual DK-halves of the catalytic component. Arrow points towards C-terminus of D-subunit, which is extended in Fe nitrogenase. (**e**) Structural alignment of putty-styled cartoon Mo, V, and Fe nitrogenases (same models as in (b)). Putty size is scaled by B-factor. Mo and Fe nitrogenase structures are measured by cryo-EM, whereas V nitrogenase structure derives from X-ray crystallography.

**Extended Data Fig. 5:**
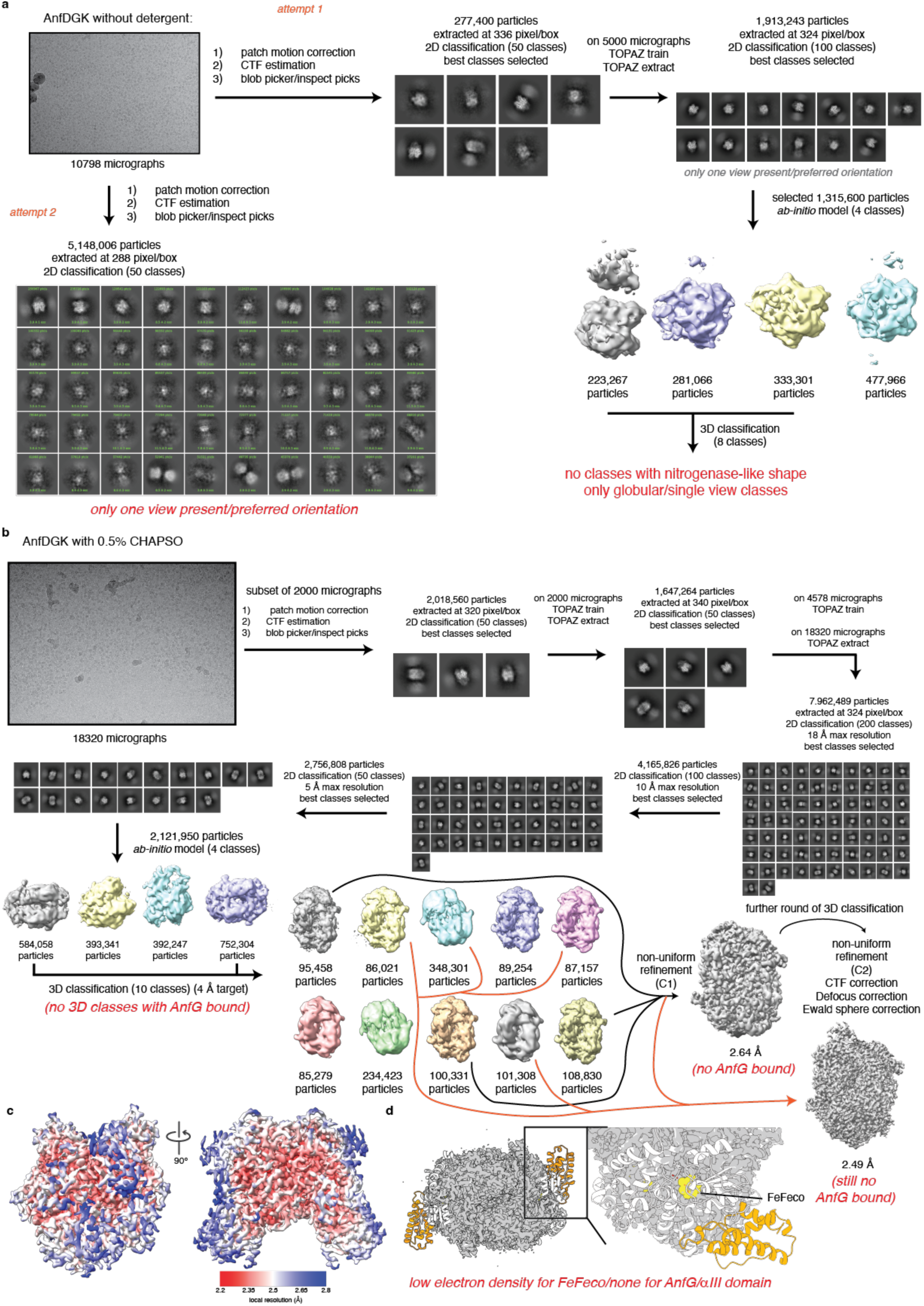
Cryogenic electron microscopy data collection and analysis of the catalytic component Anf(DGK)_2_. (**a**) Schematic data processing workflow for the electron map of Anf(DGK)_2_ collected without detergent. Dataset was collected on a Titan Krios G3i electron microscope operated at an acceleration voltage of 300 kV and equipped with a BioQuantum energy filter and a K3 direct electron detector. Dataset was processed entirely in CryoSPARC [60]. Writing in red highlights failures observed during data processing. (**b**) Schematic data processing workflow for the electron map of Anf(DGK)_2_ collected with 0.5% CHAPSO during vitrification. Dataset was collected and processed as described in (**a**). Writing in red highlights failures observed during data processing. (**c**) Local resolution as calculated by CryoSPARC mapped onto the refined density with different views shown. (**d**) 2.49 Å electron map overlaid with catalytic core of Fe nitrogenase structure (PDB: 8OIE, Anf(DGK)_2_). Zoomed view highlights lack of-or weak density surrounding AnfG subunits, α-III domains, and FeFecos. Red writing indicates findings during processing/analysis.

**Extended Data Fig. 6:**
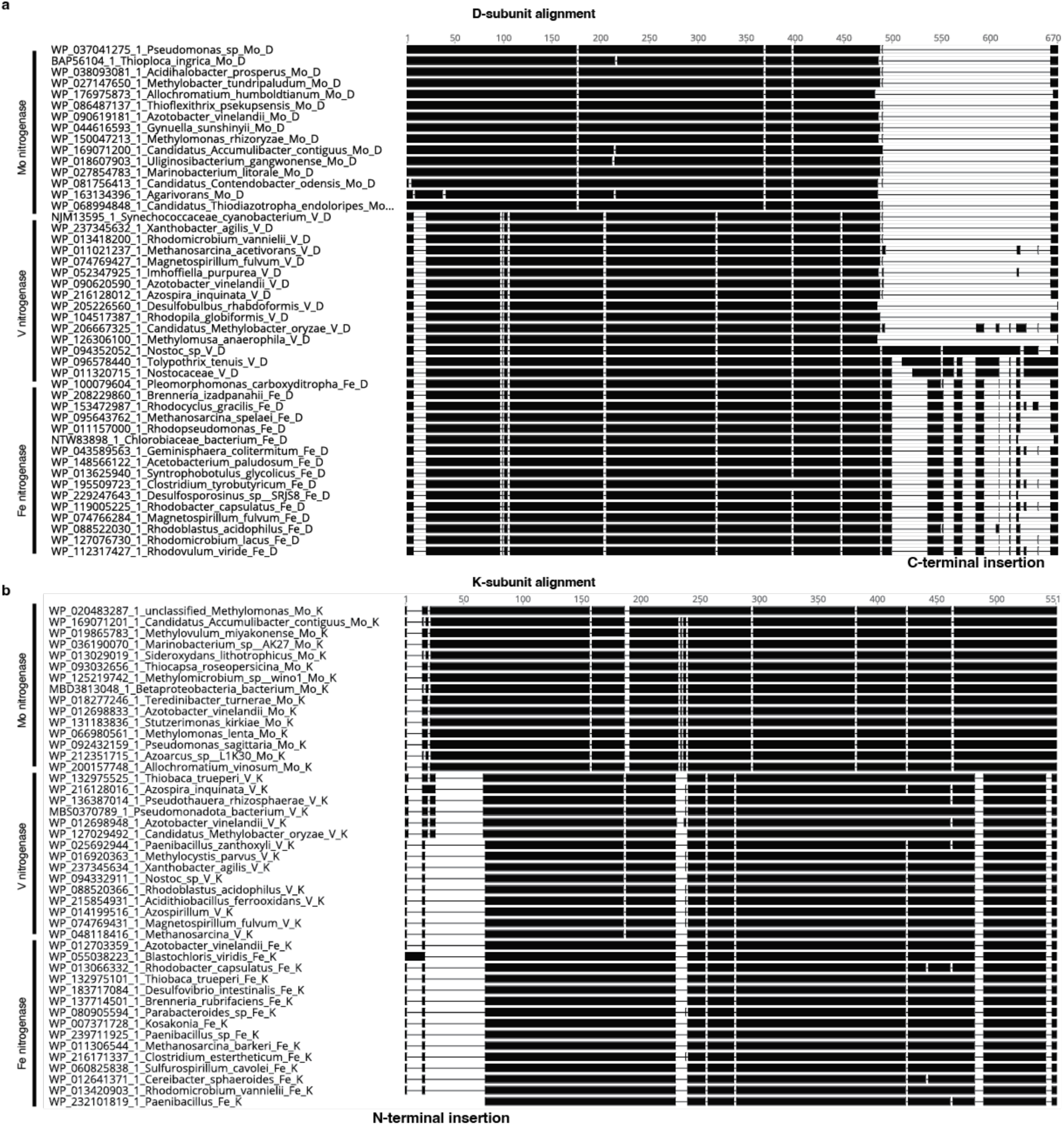
Alignments of nitrogenase D and K subunits across species. (**a**) Alignment of 46 D-subunits of representative sequence clusters (15 Mo nitrogenases, 15 V nitrogenases, and 16 Fe nitrogenases). Presence/absence of amino acids is shown in black and white, respectively. C-terminal insertion in Fe nitrogenase and select V nitrogenase sequences (cyanobacterial V nitrogenases) is highlighted. C-terminal insertion in cyanobacterial V nitrogenases is divergent of that from Fe nitrogenase in length and sequence. Amino acid sequences were aligned using MUSCLE v5 [66]. (**b**) Alignment of 45 K-subunits of representative sequence clusters (15 Mo nitrogenases, 15 V nitrogenases, and 16 Fe nitrogenases). Presence/absence of amino acids is shown in black and white, respectively. N-terminal insertion in Mo nitrogenases is highlighted. Amino acid sequences were aligned using MUSCLE v5.

**Extended Data Fig. 7:**
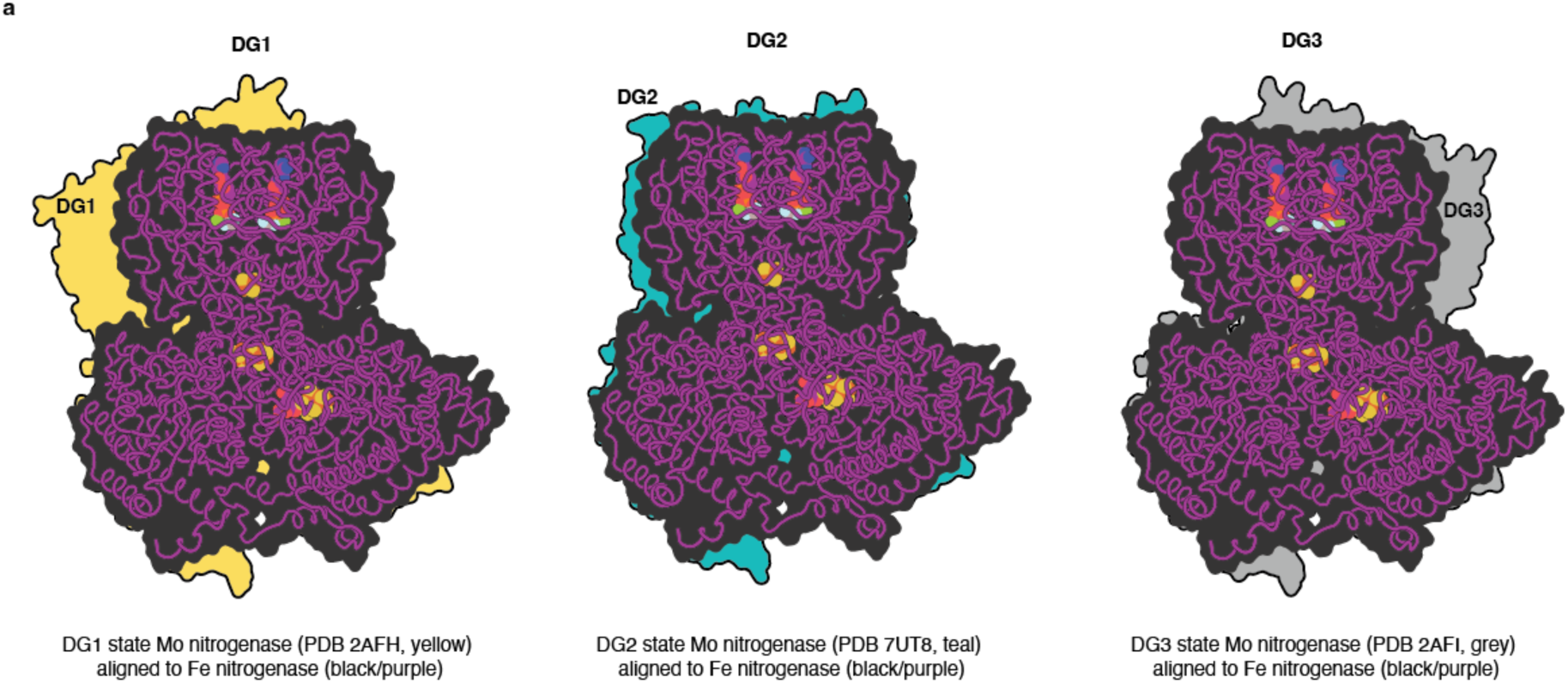
Fe nitrogenase structure docking geometry. Fe nitrogenase AnfDGKH_2_ (PDB: 8OIE) overlaid with surface outlines of nitrogenase structures containing varying reductase:catalytic component docking geometries (DG). DG1 in yellow (PDB ID: 2AFH), DG2 in teal (PDB ID: 7UT8), and DG3 in grey (PDB ID: 2AFI). Fe nitrogenase surface outline is shown in black, cartoon is shown in purple, and cofactors are shown as spheres.

