## Supplemental Tables 1 to 4 for "Structural Insights into the Iron Nitrogenase Complex"

### Supplementary Information

**Table S1: Primers used in this study.**

| Primer | Target | Sequence (5´-3´) | Purpose |
|---|---|---|---|
| oMM0021 | B10S genomic DNA | GCAGCGTGAAGCAGCCCGTTCGGAATTCCG | Construction of pMM0064 |
| oMM0023 | pRhon5Hi-2 | CTGCTTCACGCTGCCGCAAG | Construction of pMM0064 |
| oMM0027 | pBS85 | GTTTTTGGTCTCAGCCAATCCCTGGG | Construction of pMM0021 |
| oMM0028 | pBS85 | GGCTGAGACCAAAAACATATTCTCAATAAACCC | Construction of pMM0021 |
| oMM0033 | pOGG024 | CACCACAGGTCTCGGGGTTATGCAGCGGAAAAGG | Construction of pMM0002 |
| oMM0034 | pOGG024 | TCAGTAGGTCTCGAGCATGGTGAGAATCCAGGGGTCC | Construction of pMM0002 |
| oMM0035 | B10S genomic DNA | TACAACAGGTCTCGTGCTCCCGAGGCGAC | Construction of pMM0002 |
| oMM0036 | B10S genomic DNA | CATCATGGTCTCGACCGGGTAAGGAGTTCCTGTC | Construction of pMM0002 |
| oMM0037 | B10S genomic DNA | CACCACAGGTCTCGACCCATTCCAAGGGCCG | Construction of pMM0002 |
| oMM0038 | B10S genomic DNA | CAATCAGGTCTCGTGGCTGCAGATCCAGTCCGTA | Construction of pMM0002 |
| oMM0054 | B10S genomic DNA | CGTAGTCTGACGATGCGCACTTC | Construction of pMM0057 |
| oMM0055 | B10S genomic DNA | GTCAAAGGAGGCAAGCCCCATCACGAG | Construction of pMM0057 |
| oMM0056 | B10S genomic DNA | GGGCTTGCCTCCTTTGACTTGCTCGGGTT | Construction of pMM0057 |
| oMM0057 | B10S genomic DNA | GGATCCTCTAGTGCAGAACCGAATCCGAAAGC | Construction of pMM0057 |
| oMM0145 | pRhon5Hi-2 | ATCGCCGCGCAACCTGGATCCGAATTCGAGCTCCGTCGACAAGCTTG | Construction of pMM0064 |
| oMM0146 | B10S genomic DNA | CTCGAATTCGGATCCAGGTTGCGCGGCGATG | Construction of pMM0064 |

| | | | |
|---|---|---|---|
| oMM0161 | pMM0064 | AAAGCGGTGTCCGAGATGACCGGCCAGC | Construction of pMM0073 |
| oMM0162 | pMM0064 | CCGAGATCTTCGGGAGCGCCTGAAGC | Construction of pMM0073 |
| oMM0163 | pMM0064 | CATCTCGGACACCGCTTTCAGGAAGGC | Construction of pMM0073 |
| oMM0164 | pMM0064 | CCGAAGATCTCGGGCCGTCTCTTGGGC | Construction of pMM0073 |
| oMM0223 | pMM0119 | CGTTCAACCTGCCGCCGAATGGAGCCACCCGCAGTTCGAAAAAT | Construction of pMM0190 |
| oMM0224 | pMM0119 | TTTCGCTGATATCGGTCATCTGTCCCTTATTTTTCGAACTGCGGGTGGCT | Construction of pMM0190 |
| oMM0227 | pK18mobSacB | GATTTAGGTCTCTGTAAAACGACGGCCAGTGC | pK18mobSacB backbone amplification for Gibson assembly |
| oMM0228 | pK18mobSacB | GATTTAGGTCTCTCGTAATAGCGAAGAGGCCCG | pK18mobSacB backbone amplification for Gibson assembly |
| oMM0284 | B10S genomic DNA | GATTTAGGTCTCTTTACGCGCGACATCATCTTCGG | Construction of pMM0105 |
| oMM0286 | B10S genomic DNA | GATTTAGGTCTCAGAGATCCGCGCTCAGGTGC | Construction of pMM0105 |
| oMM0287 | B10S genomic DNA | GATTTAGGTCTCTTACGAACGAATATCTGGCGGCGG | Construction of pMM0105 |
| oMM0288 | B10S genomic DNA | GATTTAGGTCTCTTCTCCAGATTGGTGGAATGGCCAAGG | Construction of pMM0105 |
| oMM0323 | B10S genomic DNA | ATGCAAGCTTGGCACTGGCCGTCGTTTTACGGTCCTCGCAGCGGTC | Construction of pMM0133 |
| oMM0324 | B10S genomic DNA | CCCTAGCCCCCCCTGGAGACGAACACGCGCGA | Construction of pMM0133 |
| oMM0325 | B10S genomic DNA | GCGCGTGTTCGTCTCCAGGGGGGGCTAGGGTG | Construction of pMM0133 |

| | | | |
|---|---|---|---|
| oMM0326 | B10S genomic DNA | GCGATCGGTGCGGGCCTCTTCGCTATTACGCTCGGGTTTTCGGTGGTGG | Construction of pMM0133 |
| oMM0384 | pRhon5Hi-2 | GCCGTGGGTCGATGTTTGATGTTAGTCTTATCTGAAAGTTGTGC | Construction of pMM0119 |
| oMM0385 | pRhon5Hi-2 | CGATCTCGGCTTGAACGAATTGACCTTTTCTCCGACGAATAGAGT | Construction of pMM0119 |
| oMM0386 | pOGG024 | AATTCGTTCAAGCCGAGATCG | Construction of pMM0119 |
| oMM0387 | pOGG024 | TAACATCAAACATCGACCCACG | Construction of pMM0119 |
| oMM0389 | pMM0073 | GATTTAGGTCTCAGGAGCAGCCCGTTCGGAATTCC | Construction of pMM0119 |
| oMM0390 | pMM0073 | GATTTAGGTCTCTAGCGGGTCACCACACGTTGAGG | Construction of pMM0119 |
| oMM0394 | B10S genomic DNA | GATTTAGGTCTCTCTGGTCAAGATCCTCGACGC | Construction of pMM0131 |
| oMM0395 | B10S genomic DNA | GATTTAGGTCTCTCCAGCGACTTGCCGATACCACCTTTG | Construction of pMM0131 |
| oMM0396 | B10S genomic DNA | GATTTAGGTCTCAGGAGATCCCGATCCCGTTCGAATG | Construction of pMM0131 |
| oMM0397 | B10S genomic DNA | CAAATCGACGACCACCATGGCCAACACGCTTTTCG | Construction of pMM0131 |
| oMM0510 | pMM0119 | CCTTCCCAAGGAGCGACCCATGCACCACCATCATCACCATA | Construction of pMM0190 |
| oMM0511 | pMM0119 | CCGTAAATGGCGATCTTGCGGGTATGGTGATGATGGTGGTGC | Construction of pMM0190 |

**Table S2: Plasmids used in this study.**

| Plasmid | Code | Relevant Features | Source or Reference |
|---|---|---|---|
| pK18mobSacB | pMM0056 | Suicide vector, *oriT* (mobilizable), *sacB,* $Kan^R$ | [1] |
| pK18mobSacB-*draTG* | pMM0105 | Suicide vector, *oriT* (mobilizable), *sacB,* $Kan^R$, homologous recombination sites for draTG locus | This study |
| pK18mobSacB-*gtaI* | pMM0133 | Suicide vector, *oriT* (mobilizable), *sacB,* $Kan^R$, homologous recombination sites for gtaI locus | This study |

| | | | |
|---|---|---|---|
| pK18mobSacB-*nifHDK* | pMM0131 | Suicide vector, *oriT* (mobilizable), *sacB,* Kan$^R$, homologous recombination sites for nifHDK locus | This study |
| pK18mobSacB-*modABC* | pMM0057 | Suicide vector, *oriT* (mobilizable), *sacB,* Kan$^R$, homologous recombination sites for modABC locus | This study |
| pBS85 | pMM0017 | Suicide vector, *oriT* (mobilizable), Tet$^R$ | [2] |
| pBS85-BsaI | pMM0021 | Suicide vector, *oriT* (mobilizable), Tet$^R$, BsaI cutting site | This study |
| pBS85-BsaI-*genR* | pMM0002 | Suicide vector, *oriT* (mobilizable), Tet$^R$, BsaI cutting site, Gen$^R$, homologous recombination sites for anfHDGK locus | This study |
| pRhon5Hi-2 | pMM0031 | Broad host range plasmid, Kan$^R$ | [3] |
| pRhon5Hi-2-*anfHDGK* | pMM0064 | Broad host range plasmid, *anfHDGK* operon, Kan$^R$ | This study |
| pRhon5Hi-2-*anfHDGK*-Golden Gate | pMM0073 | Broad host range plasmid, *anfHDGK* operon with silent mutation in BsaI cutting site, Kan$^R$ | This study |
| pOGG024 | pMM0114 | Broad host range plasmid, *oriT* (mobilizable), *lacZα* cassette for golden gate cloning (BsaI), Gen$^R$ | Addgene plasmid #113991 |
| pOGG024-*kanR* | pMM0119 | Broad host range plasmid, *oriT* (mobilizable), *lacZα* cassette for golden gate cloning (BsaI), Kan$^R$ | This study |
| pOGG024-*kanR*-*anfHDGK* | pMM0119 | Broad host range plasmid, *oriT* (mobilizable), *anfHDGK* operon, Kan$^R$ | This study |
| pOGG024-*kanR*-*anfHDGK*-Strep/His | pMM0190 | Broad host range plasmid, *oriT* (mobilizable), *anfHDGK* operon with N-terminal His$_6$-tag and C-terminal Strep tag II, Kan$^R$ | This study |

**Table S3: Strains used in this study.**

| Strain | Genotype | Source or Reference |
|---|---|---|
| *Rhodobacter capsulatus* B10S | Wildtype | [4] |
| *Rhodobacter capsulatus* MM0425 | Δ*anfHDGK*::*genR* Δ*modABC* Δ*draTG* Δ*gtaI* Δ*nifHDK* | This study |
| *Rhodobacter capsulatus* MM0436 (expression strain) | Δ*anfHDGK*::*genR* Δ*modABC* Δ*draTG* Δ*gtaI* Δ*nifHDK* | This study |
| *Escherichia coli* DH5α | $F^-$ Φ80*lac*ZΔM15 Δ(*lac*ZYA-*arg*F) U169 *rec*A1 *end*A1 *hsd*R17($r_K^-$, $m_K^+$) *pho*A *sup*E44 *thi*-1 *gyr*A96 *rel*A1 $\lambda^-$ | Thermo Fisher Scientific Inc. (Waltham, USA) catalogue #18265017 |
| *Escherichia coli* ST18 | RP4-2 *Tc*::*Mu* *Km*::*Tn7* Δ*hemA* mutant | [5] |

**Table S4: CryoEM data collection, refinement and model statistics.**

| | |
|---|---|
| **Data collection** | |
| Microscope | Titan Krios G3i |
| Voltage (kV) | 300 |
| Camera | K3 |
| Magnification | 105,000 |
| Pixel size at detector (Å/pixel) | 0.837 |
| Total electron exposure ($e^-$/Å$^2$) | 50 |
| Exposure rate ($e^-$/pixel/sec) | 15 |
| Number of frames collected during exposure | 50 |
| Defocus range (μm) | -1.4 to -2.4 |
| Automation software | EPU, CryoSPARC, TOPAZ |
| Energy filter slit width | -30 eV |
| Micrographs collected (no.) | 10148 |
| Micrographs used (no.) | 4601 |
| Total extracted particles (no.) | 3,014,316 |
| Refined particles (no.) | 432,216 |
| Final particles (no.) | 218,653 |
| Point-group or helical symmetry parameters | C 2 |
| Resolution (global, Å) | 2.35 |
| $FSC_{0.143}$ (unmasked / masked) | 3.00 / 2.35 |
| Resolution range (local, Å) | 1.83 - 7.30 |
| Map sharpening B-factor (Å$^2$) / (B-factor Range) | -50.0 |
| Map sharpening methods | CryoSparc sharpening |
| **Model composition** | |
| Protein residues | 3262 |
| Ligands | 4 × ADP, 4 × $AlF_3$, 4 × $Mg^{2+}$, 2 × [$Fe_8S_7$], 2 × [$Fe_8S_9C$], 2 × (*R*)-homocitrate, 2 × [$Fe_4S_4$] |
| **Model Refinement** | |
| Refinement package | Phenix |
| - method | real space |
| - resolution cutoff | 2.34 |

| | |
|---|---|
| Model-Map scores | |
| - $CC_{Map}$ | 0.82 |
| - Average $FSC_{0.143}$ (unmasked / masked) | 2.34 / 2.16 |
| B factors ($Å^2$) | |
| Protein residues (min / max / mean) | 10.50 / 116.09 / 61.09 |
| Ligands (min / max / mean)<br>Waters (min / max / mean) | 34.62 / 93.59 / 66.10<br>34.72 / 75.14 / 53.52 |
| R.M.S. deviations from ideal values | |
| Bond lengths (Å) | 0.012 |
| Bond angles (°) | 0.845 |
| **Validation** | |
| MolProbity score | 1.42 |
| CaBLAM outliers | 0.93 |
| Clashscore | 5.12 |
| Poor rotamers (%) | 0.44 |
| C-beta deviations | 0.00 |
| EMRinger score | 5.26 |
| Ramachandran plot | |
| Favored (%) | 97.16 |
| Outliers (%) | 0.00 |
